# Evaluating the performance of splicing predictors on thousands of synthetic gene variants

**DOI:** 10.64898/2026.08.24.746734

**Authors:** Fernando Bellido Molias, Grzegorz Kudla

## Abstract

Computational predictors of RNA splicing are increasingly used to interpret genetic variants and to design synthetic genes, yet they are almost always benchmarked on endogenous human sequences closely related to their training data. Whether their performance reflects genuine recognition of splicing signals, or instead exploits statistical features of natural genomes such as conservation and exon–intron composition, remains unclear. Here we benchmark eleven splicing predictors on thousands of synthetic GFP variants that are heavily recoded and dissimilar from any training data, using long-read sequencing to measure splicing directly at each position. Despite this distribution shift, modern deep-learning predictors retained strong performance, and the resulting ranking was largely stable across position-level and construct-level benchmarks. SpliceTransformer ranked highest, followed by AlphaGenome and SpliceAI. Tools that ignore long-range sequence context performed substantially worse, largely because they assign high scores to many non-spliced positions. This ranking broadly agrees with benchmarks on endogenous variants, indicating that the leading models capture transferable, sequence-intrinsic determinants of splicing. We further provide a unified calibration that maps each predictor’s scores onto the measured fraction of spliced reads, allowing scores to be interpreted as splicing outcomes and compared directly between tools. Our results show that current deep-learning models generalise beyond natural genomes and provide a practical framework for splicing-aware sequence design.

## Introduction

Recent years have seen large advances in computational tools that take a biological sequence as their input and return a predicted functional outcome such as protein structure [1], promoter activity [2], variant pathogenicity [3], or gene regulatory information [4]. Compared to other genotype–phenotype prediction settings, splicing poses unique challenges. Splicing depends not only on local elements such as donor and acceptor sites, but also on the relative location of these elements within the sequence and on the action of RNA-binding proteins that interact with distal sequence elements in a context-specific manner [5, 6]. Experimental studies have further shown that the effect of a splice-regulatory mutation can depend strongly on the strength of neighbouring splice sites and surrounding regulatory elements, highlighting the importance of sequence context in splice-site recognition [7]. Because of this, splicing prediction was long considered particularly difficult.

Splicing predictors have evolved from local models that score individual sites in isolation toward methods that draw on progressively larger sequence contexts to capture the dependencies that govern splice-site selection (Table 1) [4, 8–17]. Early motif- and statistical-based approaches [8, 9] evaluate splice sites through local sequence features and therefore suffer from high false-positive rates when broader sequence context is not considered. NNSplice [10] and GeneSplicer [11] introduced machine learning to improve the modeling of these local sequence determinants, while SpliceFinder [12] and Spliceator [13] increased representation capacity through the use of convolutional neural networks. More recently, the integration of deep-learning architectures with increasingly large sequence contexts has substantially advanced the accuracy of splicing predictions. SpliceAI [14], a deep convolutional neural network, enabled accurate prediction of both canonical and cryptic splice sites from primary sequence using large sequence contexts; Pangolin [15] extended this paradigm by incorporating tissue-aware modeling of splice-site usage; SpliceTransformer [16] introduced an attention-based architecture to capture long-range dependencies; and OpenSpliceAI [17] provided an open-source reimplementation of SpliceAI, enabling custom retraining. Most recently, AlphaGenome [4] introduced a foundation-model approach, jointly predicting splicing and other regulatory outputs across very large genomic windows.

**Table 1.**
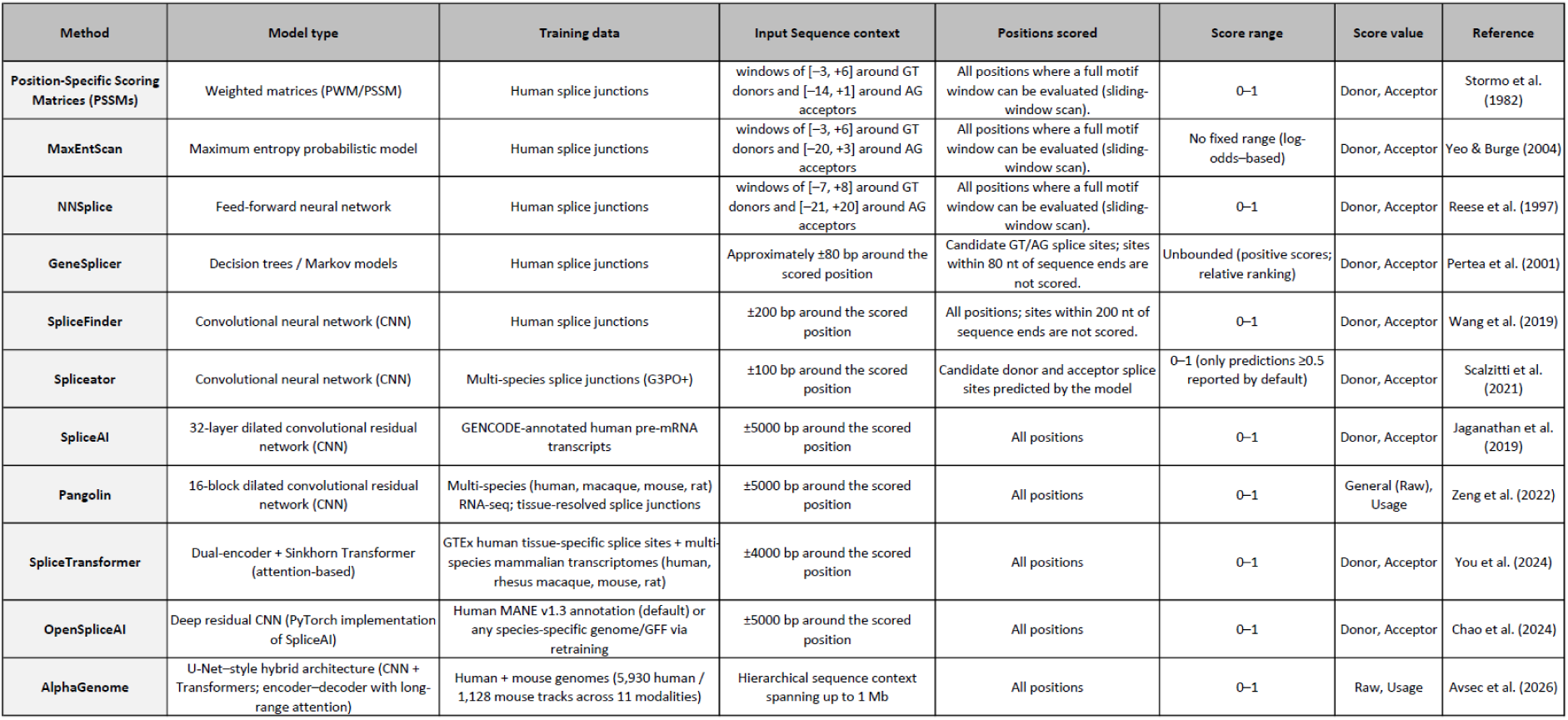
Characteristics of splice-site prediction methods included in the benchmark. The table summarizes the architecture, training data, sequence context, evaluated positions, score range, output type, and original publication for each splice-site prediction method analysed in this study. Sequence context denotes the amount of surrounding sequence used by each model when generating predictions. Evaluated positions indicate whether scores are reported for all sequence positions or only for candidate splice sites identified by the method.

Splicing predictors are commonly trained on sets of endogenous human genes and evaluated on another set of genes (e.g. from a different chromosome) that has not been used for training. While important, this benchmark risks data leakage due to similarities between genes used for training and testing. In addition, systematic differences in nucleotide composition between exons and introns could allow predictors to rely on local GC content, codon usage, and similar properties as a proxy for exon–intron boundaries [18]. As a result, strong performance on unseen endogenous genes does not necessarily indicate that the model has learned the splicing grammar. Another common benchmarking approach relies on discriminating between benign and pathogenic mutations [19] and predicting the outcomes of massively parallel reporter assays (MPRAs) [20]. High performance of modern splice predictors in these tasks indicates that they are generally suitable for the interpretation of naturally occurring genetic variants. However, most of these tasks represent in-distribution predictions, because the variants predicted typically only differ by a single nucleotide from the training data. AI tools are known to perform better within the training distribution, and performance can drop substantially outside of it. It is also possible that predictors rely on sequence conservation as an indicator of which positions are functionally important. As a result, it is not clear whether strong performance on existing benchmarks generalizes to out-of-distribution predictions for genuinely novel genes.

This question is particularly relevant for applications in basic research, bioengineering, and gene therapy. In these contexts, constructs are often assembled from fragments of unrelated genes, and are frequently tagged or codon-optimized. This can introduce unintended splice-regulatory elements, leading to aberrant splicing. Indeed, aberrant RNA splicing has been repeatedly observed in plasmid-expressed cDNAs across diverse genes despite the absence of annotated introns, including *ABCB1* [21], *ACE2* and *ALKBH5* [22], *CAPN10* [23], *CCR5* [24], *DUX4* [25], *EIF4G* [26], *ERCC1* [27], *FACI* [28], *HBG1* and *HBG2* [29], *IL1RN* [30], *MYC* [31], *NRF1* [26], *RBM3* [26] and *XIAP* [26, 31, 32], among others. Mechanistic studies indicate that these effects frequently arise from activation of cryptic splice sites within the insert(s) or the plasmid backbone, generating unintended transcript isoforms. These findings demonstrate that even wild-type sequences derived from fully processed mRNAs remain susceptible to reprocessing by the host splicing machinery in non-native contexts. Consistent with these observations, alterations in splicing have also been reported as a consequence of codon optimization. In SARS-CoV-2 constructs, sequence optimization introduced functional splice sites that generated unintended transcript isoforms and protein products absent from the wild-type sequence, highlighting that synonymous recoding can give rise to novel, non-native protein species with potentially clinically relevant consequences [33, 34]. We recently found this to be pervasive rather than anecdotal: across thousands of synonymous GFP variants and hundreds of intronless human ORFs, roughly half of constructs showed detectable splicing [35]. Such events are easily missed by short-read sequencing, which often fails to resolve full-length isoforms, but are captured directly by long-read sequencing.

Here, we directly test whether the strong benchmark performance of current predictors extends to sequences unlike those they were trained on. We leverage our data from thousands of synthetic GFP variants to benchmark eleven splicing predictors. These variants span a wide range of nucleotide composition and codon usage bias, were expressed in human cells, and their transcript structures were resolved by long-read cDNA sequencing, providing direct ground truth for splicing events and junction positions. We evaluate predictors at three levels: position-level detection of cryptic splice sites, construct-level splicing propensity, and the relationship between prediction scores and the thresholds needed to prioritize sequences in practice. Because these sequences are distant from any training data and free of the confounders present in natural genomes, the benchmark isolates sequence-intrinsic splice-site recognition, providing a stringent evaluation of whether current tools have learned generalizable splicing rules.

## Results

### A large synonymous GFP library enables systematic benchmarking of splicing predictors

To enable benchmarking of splicing predictors, we employed our synonymously recoded GFP library, in which extensive nucleotide diversity was introduced by randomizing the third position of each codon across the coding sequence [35]. Degenerate codons were biased toward GC-rich nucleotides and high codon adaptation (average GC3 ∼84% and mean CAI ∼0.75) to reflect sequence composition common among natural human genes and transgenes used in biomedical applications. The resulting library comprises extensive synonymous variation (average pairwise Hamming distance, ∼85 nucleotides), representing a diverse set of genes not seen by current splicing predictors. Each construct was cloned into a barcoded mammalian expression vector, enabling pooled functional assays and unambiguous assignment of sequencing reads to individual variants (Fig. 1A).

**Figure 1.**
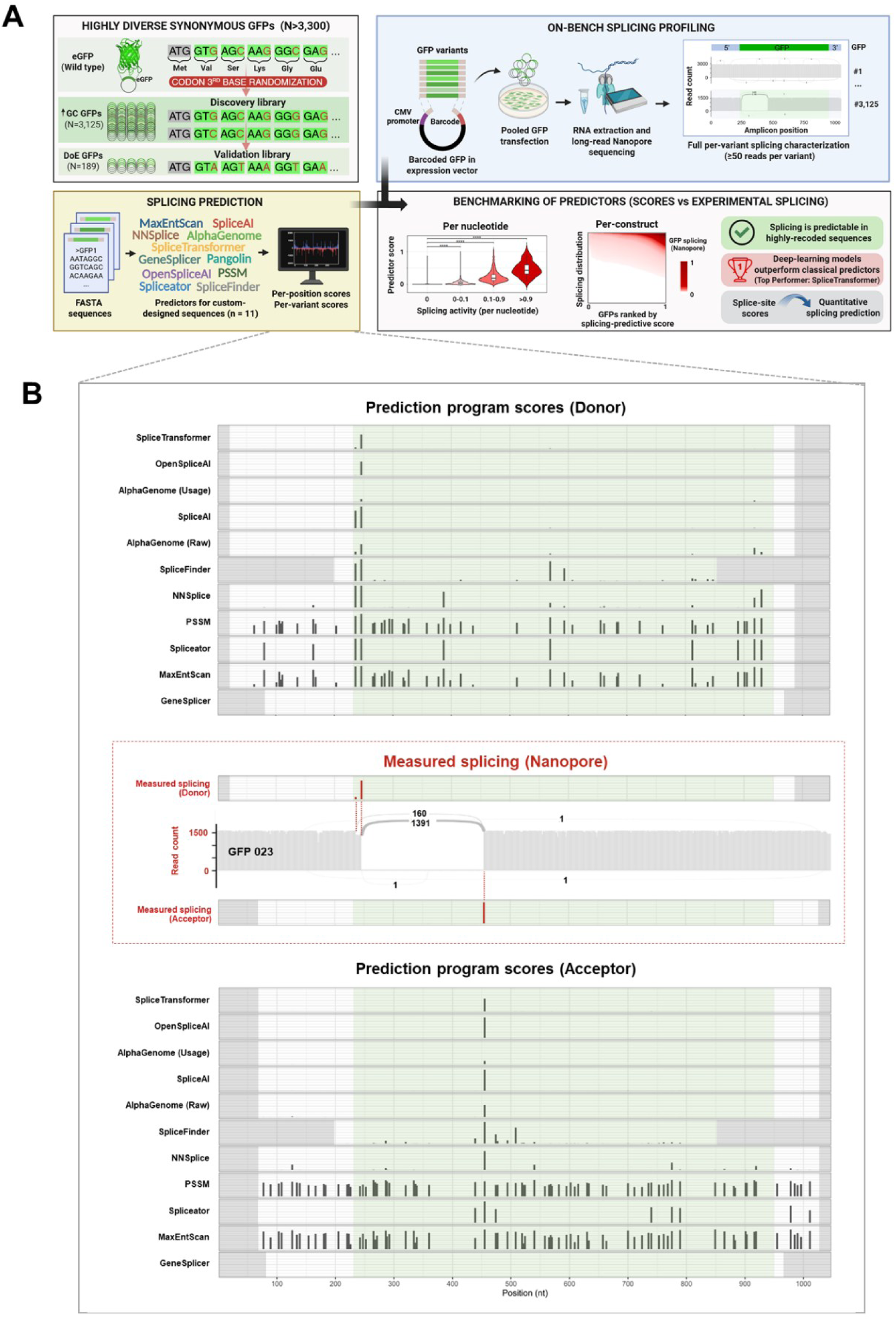
Workflow for benchmarking splicing prediction in synthetic constructs. **(A)** Overview of the benchmarking workflow. A synonymous GFP library was generated by third-codon position randomization, producing >3,000 variants with extensive synonymous diversity. Each construct was cloned into a barcoded mammalian expression vector, transiently expressed in human cells, and profiled by long-read Nanopore sequencing to quantify full-length transcript isoforms and nucleotide-resolution splice-site usage. In parallel, splice-site scores were computed across each sequence using the indicated predictors. Predictor performance was benchmarked at the nucleotide level (splice-site usage per position) and construct level (overall splicing propensity). **(B)** Representative example of splice-site prediction across a synthetic GFP construct. Predicted donor and acceptor scores generated by splicing predictors are compared with splice-site usage experimentally determined by long-read Nanopore sequencing.

To characterize sequence-specific splicing outcomes across the library, we re-analysed transcriptomic data from pooled constructs transiently transfected into HeLa cells, followed by RNA extraction, reverse transcription, and long-read Oxford Nanopore sequencing. Applying a stringent coverage threshold (≥50 UMI-corrected reads per construct; UMI = unique molecular identifier), we retained 3,125 GFP variants with high-confidence transcript profiles, revealing hundreds of distinct splicing isoforms arising from activation of cryptic splice sites within the coding sequence. Cryptic splicing was widespread across the library, with 64.9% of variants showing detectable splicing. On average, 9.3% of transcripts per construct were spliced (95% CI: 8.6–10.0%). For every nucleotide position in each construct, we quantified donor and acceptor splice-site usage, defined as the fraction of UMI-corrected reads in which that position participated in a splice junction. These data provide a high-resolution, systematic map of cryptic splice-site usage within a controlled synthetic coding context.

### Splicing predictors capture cryptic splice signals in synthetic coding sequences

We assessed splice-site prediction in these synthetic constructs across eleven widely used predictors spanning motif-based and machine learning architectures (Table 1). Predicted donor and acceptor scores were compared with experimentally measured splice-site usage at each position. Because ∼99% of splice junctions in human transcriptomes occur at canonical splice-site dinucleotides [36], analyses were restricted to positions compatible with GT (donor) and AG (acceptor) motifs. This ensured that performance comparisons were conducted only among physiologically plausible candidate sites, avoiding inflation of predictive accuracy caused by the large number of non-spliceable positions. An example of this analysis for a representative GFP variant is shown in Fig. 1B: the majority of predictors assign elevated scores to the experimentally observed splice sites. However, differences in scoring strategies across methods lead to substantial variation in predicted outcomes, motivating the systematic benchmarking of splicing predictors.

To examine how predicted scores relate to experimentally observed splicing activity, GT/AG positions were stratified into ten bins (deciles) of splice-site usage, ranging from null (no detected splicing) to very high (>0.9). Across most predictors, higher splice-site usage was associated with progressively higher predicted scores (Fig. 2A). Deep learning models such as SpliceTransformer, AlphaGenome, and SpliceAI showed markedly improved separation between usage categories, with predicted scores increasing consistently with observed splicing activity. Classical motif-based predictors, including MaxEntScan, NNSplice, and PSSM models, showed separation between unspliced (splice-site usage=0) and spliced (>0) positions, but struggled with differentiating between low- and high-usage splice sites.

**Figure 2.**
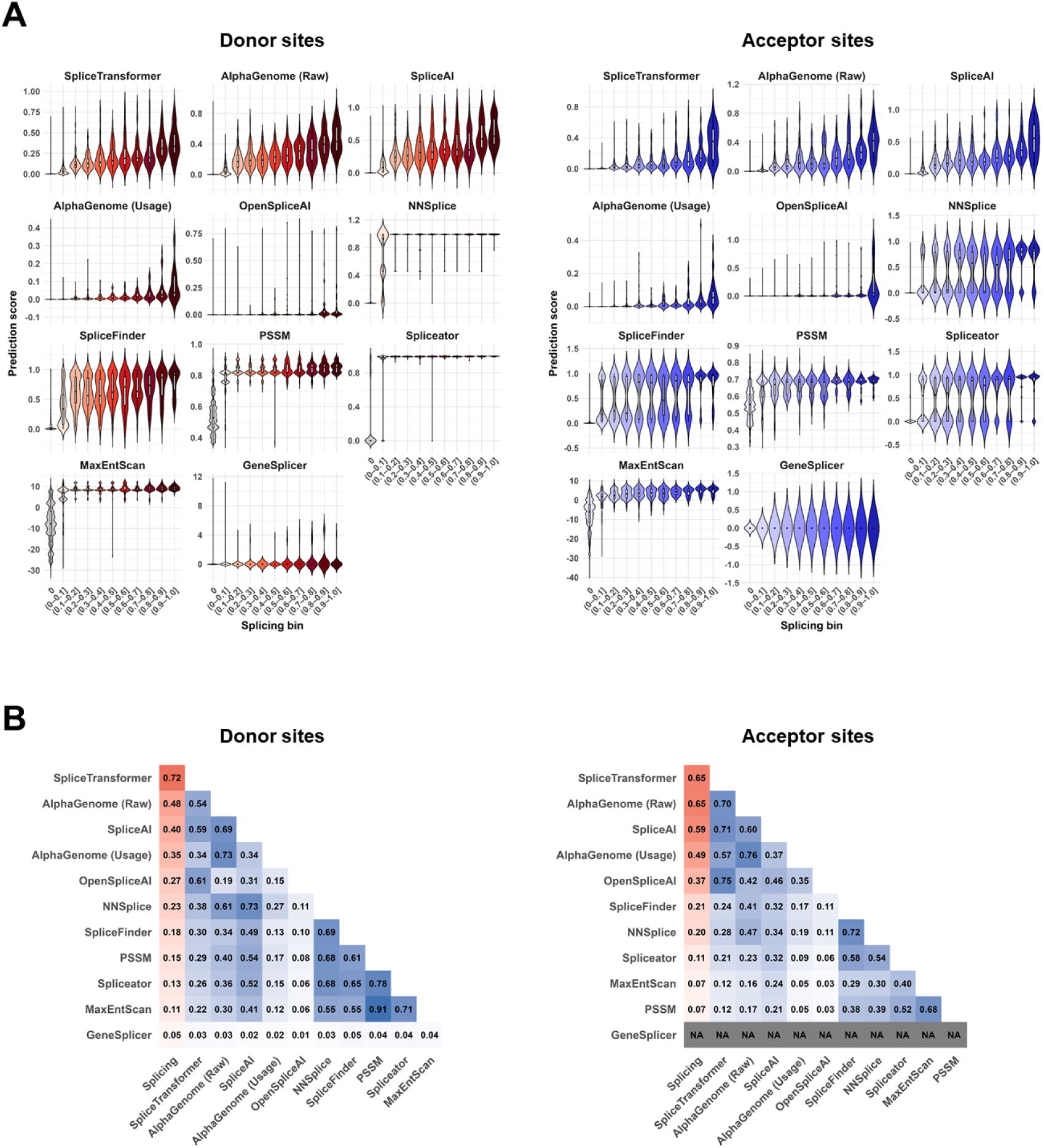
Deep learning predictors show high agreement with experimentally measured splice-site usage in synthetic constructs. **(A)** Distributions of per-site predictor scores across 3,125 GFP variants grouped according to experimentally measured splice-site usage derived from long-read Nanopore sequencing. Data are shown for splicing-compatible positions (GT donor sites and AG acceptor sites). **(B)** Pearson correlations between predicted scores and experimentally measured splice-site usage (red column) and pairwise correlations between predicted scores (blue columns).

To quantify the agreement between predictors and experimental measurements, we computed pairwise Pearson correlations between predicted scores and observed splice usage across all compatible positions (Fig. 2B). Across donor sites, SpliceTransformer (r = 0.72), AlphaGenome raw scores (r = 0.48), and SpliceAI (r = 0.40) showed the strongest correlations with measured splice usage. Similar trends were observed for acceptor sites, where SpliceTransformer (r = 0.65), AlphaGenome raw scores (r = 0.65), and SpliceAI (r = 0.59) again showed the highest agreement. These top-performing predictors integrate extended sequence context through deep neural architectures. In contrast, motif-based predictors fail to capture combinations of weaker regulatory signals distributed across broader sequence contexts.

When correlating prediction methods with each other (Fig. 2B), distinct groupings emerged reflecting underlying modeling approaches. MaxEntScan and PSSM exhibited the highest mutual correlation among donor predictors (r ≈ 0.91), consistent with their shared reliance on local sequence motif scoring. Deep learning models — including SpliceTransformer, AlphaGenome, SpliceAI, and OpenSpliceAI — formed a distinct cluster, showing stronger correlations with experimental splice usage and with each other, in line with their use of extended sequence context and deep neural architectures.

To test whether these observations generalize beyond GC-rich synonymous constructs, we analyzed an independent GFP library (n = 189) generated through a factorial design-of-experiments (DoE) framework [35]. This smaller library systematically varies key sequence properties relevant to synthetic gene design, including GC content, CpG dinucleotide frequency, and codon adaptation index (CAI), exploring a broader range of design parameters and larger sequence variation than the high-GC library (average pairwise Hamming distance ∼147 nucleotides; Suppl. Fig. 1A, Suppl. Fig. 1B). Despite these differences, performance trends across predictors remained consistent (Suppl. Fig. 1C). SpliceTransformer, AlphaGenome, and SpliceAI showed the strongest agreement with experimentally measured splicing outcomes (Suppl. Fig. 1D), while motif-based predictors showed weaker performance.

### SpliceTransformer provides the most accurate classification across splicing thresholds

The above analyses suggest that some predictors (e.g. MaxEntScan, Spliceator, NNSplice) can differentiate spliced from unspliced positions, but struggle to distinguish positions with low versus high levels of splicing (Fig. 2A). To test this systematically, we examined how well predictors classified splice sites into low- and high-spliced groups across a range of splicing-level cutoffs used to define those groups. We first quantified ROC-AUC at several splicing thresholds, between 2% and >50% (Suppl. Fig. 2A). Across predictors, performance was consistently high, particularly for the 50% splicing threshold, where several models approached near-perfect classification. Donor-site prediction showed uniformly high performance across methods (ROC-AUC typically 0.92–0.999), with SpliceTransformer achieving the strongest discrimination. In contrast, acceptor-site prediction exhibited greater variability (ROC-AUC 0.76–0.998), likely reflecting the increased regulatory complexity of acceptor recognition.

The high accuracy of all methods likely reflects the strong class imbalance inherent to splicing data, where most GT/AG sites are not used for splicing and have low scores across all predictors. To account for this, we next evaluated performance using precision–recall curves (PR-AUC), which are more informative for rare positive events. Under this framework, differences between predictors became more pronounced: SpliceTransformer showed the strongest performance for donor-site prediction across thresholds, whereas for acceptor sites, SpliceTransformer, SpliceAI, and AlphaGenome formed a top-performing group with comparable performance, consistently outperforming other predictors (Fig. 3A, Suppl. Fig. 2B). To summarize performance across the full range of splicing activity, we computed PR-AUC across a continuous range of thresholds, from null to almost complete splicing. This analysis confirmed the overall ranking, with SpliceTransformer consistently achieving the highest average performance, followed by SpliceAI and AlphaGenome (Fig. 3B). Most predictors performed best at low splicing thresholds, indicating more effective discrimination of unspliced variants.

**Figure 3.**
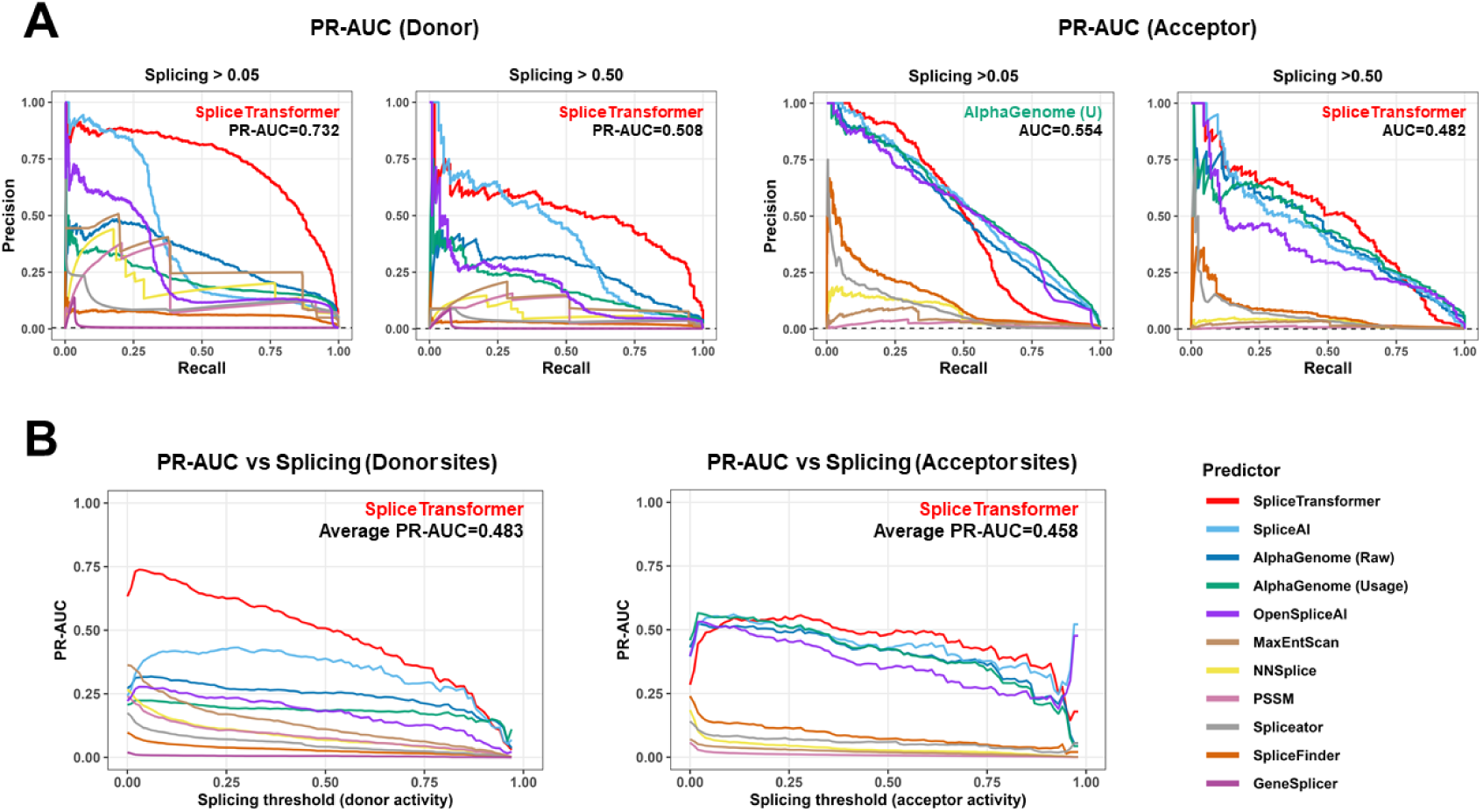
Benchmarking predictor performance across splicing activity thresholds. **(A)** Precision–recall (PR) curves for donor and acceptor splice-site classification at representative splicing thresholds corresponding to moderate (0.05) and highly active (0.50) splice-site usage. (**B**) PR-AUC values for donor and acceptor splice-site classification across a continuous range of splicing thresholds. Average PR-AUC values are indicated for the top-performing predictors.

We next repeated these analyses in the smaller design-of-experiments (DoE) library (Suppl. Fig. 3). As before, donor-sites were classified more accurately than acceptor sites. SpliceTransformer remained the best predictor for donor sites at low splicing thresholds, whereas SpliceAI showed stronger performance for highly active sites and acceptor prediction. Similar trends were observed in splice-site prioritization analyses aimed at identifying functional splice sites. Consistent with the above benchmarks, SpliceTransformer showed the strongest overall prioritization performance. For example, it identified 90% of donor sites with >10% splice-site usage among the top <1% of ranked positions (Suppl. Fig. 4). Together, these results demonstrate that modern deep learning–based predictors achieve robust and generalizable performance across diverse synthetic sequence contexts.

### Calibrating predictor scores to splicing probabilities

The above results suggest that predictor scores can serve as quantitative estimates of the fraction of spliced reads at each position. However, because different predictors are trained on different objectives and produce scores on different scales, a given score value cannot be assumed to correspond to the same expected level of splicing across methods. To calibrate scores, we modeled the probability of observing different levels of splicing as a function of predictor score using generalized additive models, which fit a smooth, flexible curve without assuming the relationship is linear. This revealed distinct calibration profiles across predictors (Fig. 4). For example, a SpliceTransformer donor score of 0.2 corresponds to a 50% probability of observing more than 30% spliced reads at that position (purple band, top left), whereas the same score from SpliceAI corresponds to only a 1.15% chance of exceeding 30% spliced reads (purple), and a 12.0% chance of observing any splicing at all (yellow, orange, red, and purple). For deep learning predictors, a high acceptor score typically indicates a very high likelihood of splice site usage, whereas high donor scores do not necessarily convert to high usage. Motif-based and early machine learning predictors do not reach high splice site usage even at high scores, indicating limited specificity (Fig. 4).

**Figure 4.**
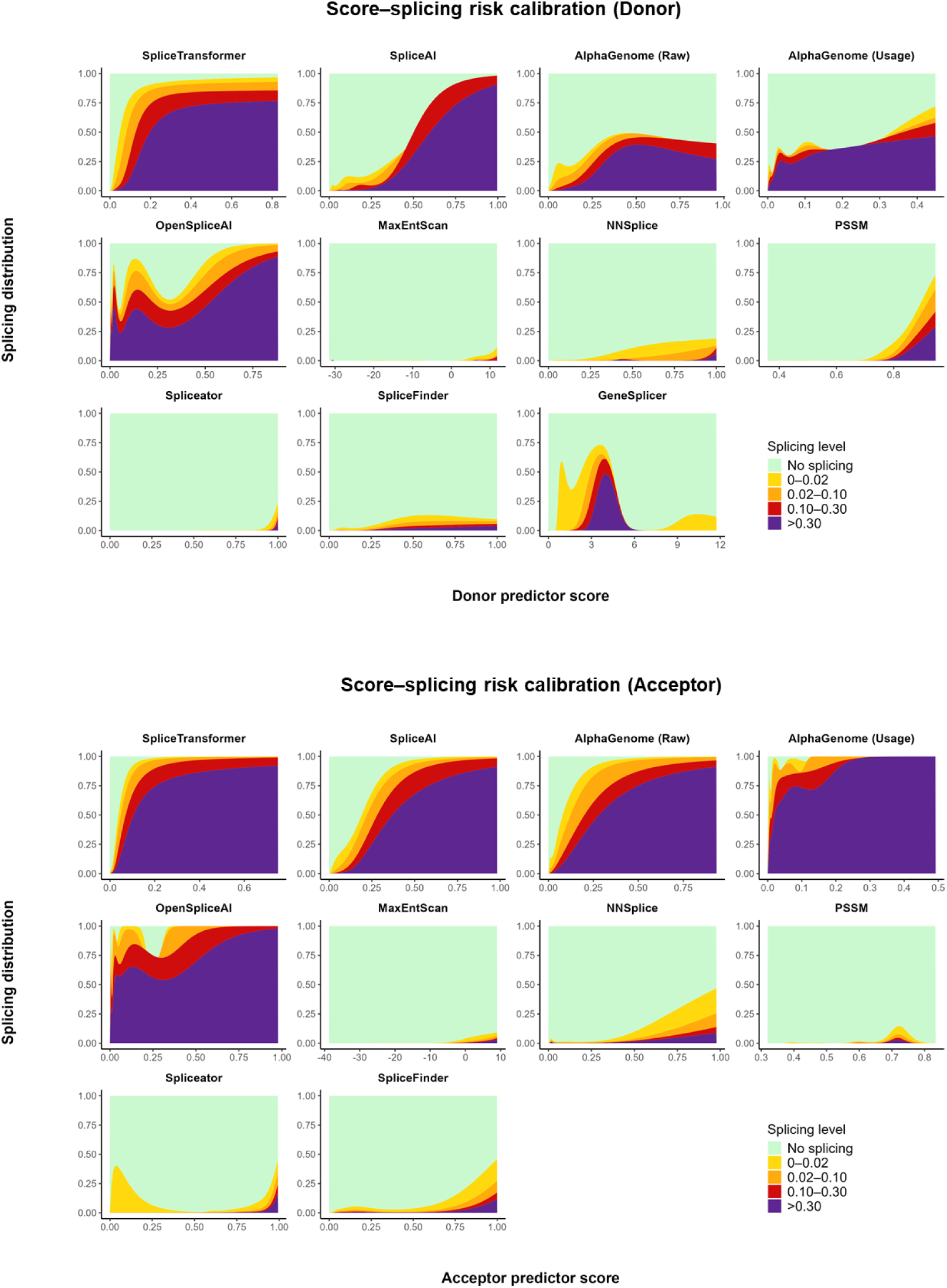
Score–splicing calibration across all evaluated predictors. Generalized additive models (GAMs) were used to estimate the probability of observing each level of splice-site activity as a function of predictor score, shown separately for donor (top) and acceptor (bottom) sites. At each score, the bands show the probability of each splicing outcome and sum to 1: the purple band at the bottom is the probability of observing >0.30 spliced reads; the red, orange and yellow bands indicate progressively smaller fractions of spliced reads; and the green band at the top is the probability of observing no spliced reads (see legend). Predictors are ordered by average PR-AUC.

To facilitate the interpretation of predictor outputs, we provide reference score thresholds for recovering different fractions of experimentally detected splice sites across distinct levels of splicing activity (Suppl. Figs. 5, 6). Top-performing predictors showed progressive enrichment of strongly spliced sites at higher scores while simultaneously stratifying splice sites according to splicing intensity, while motif-based predictors retained a substantial proportion of non-spliced positions among high-scoring predictions.

### Predicting splicing propensity at the gene level

Most predictors are designed to score individual splice junctions and do not address the pairing of donor sites with their respective acceptors. The exception is AlphaGenome, which explicitly models donor–acceptor pairing. In many applications, however — such as the design of constructs for basic research, bioengineering, or gene therapy — what matters is the overall splicing propensity of the construct rather than the exact usage of individual sites. We therefore evaluated several strategies for aggregating the full set of predicted donor and acceptor scores into a single whole-gene metric of splicing propensity (Fig. 5A). These fell into three groups: maxima-based methods (the maximum donor score, the maximum acceptor score, or the sum of the maximum donor and maximum acceptor scores), sums over all candidate sites (the summed donor, acceptor, or combined scores across the sequence), and donor–acceptor pairing schemes. For the pairing schemes, we computed the highest pair score across all compatible donor–acceptor pairs, where a pair is deemed compatible if the acceptor site is at least 50 nucleotides downstream from the donor site; donor and acceptor scores were combined either as their arithmetic sum (ArithmPaired) or their geometric mean (GeomPaired). Only GT donor and AG acceptor dinucleotides were considered throughout.

**Figure 5.**
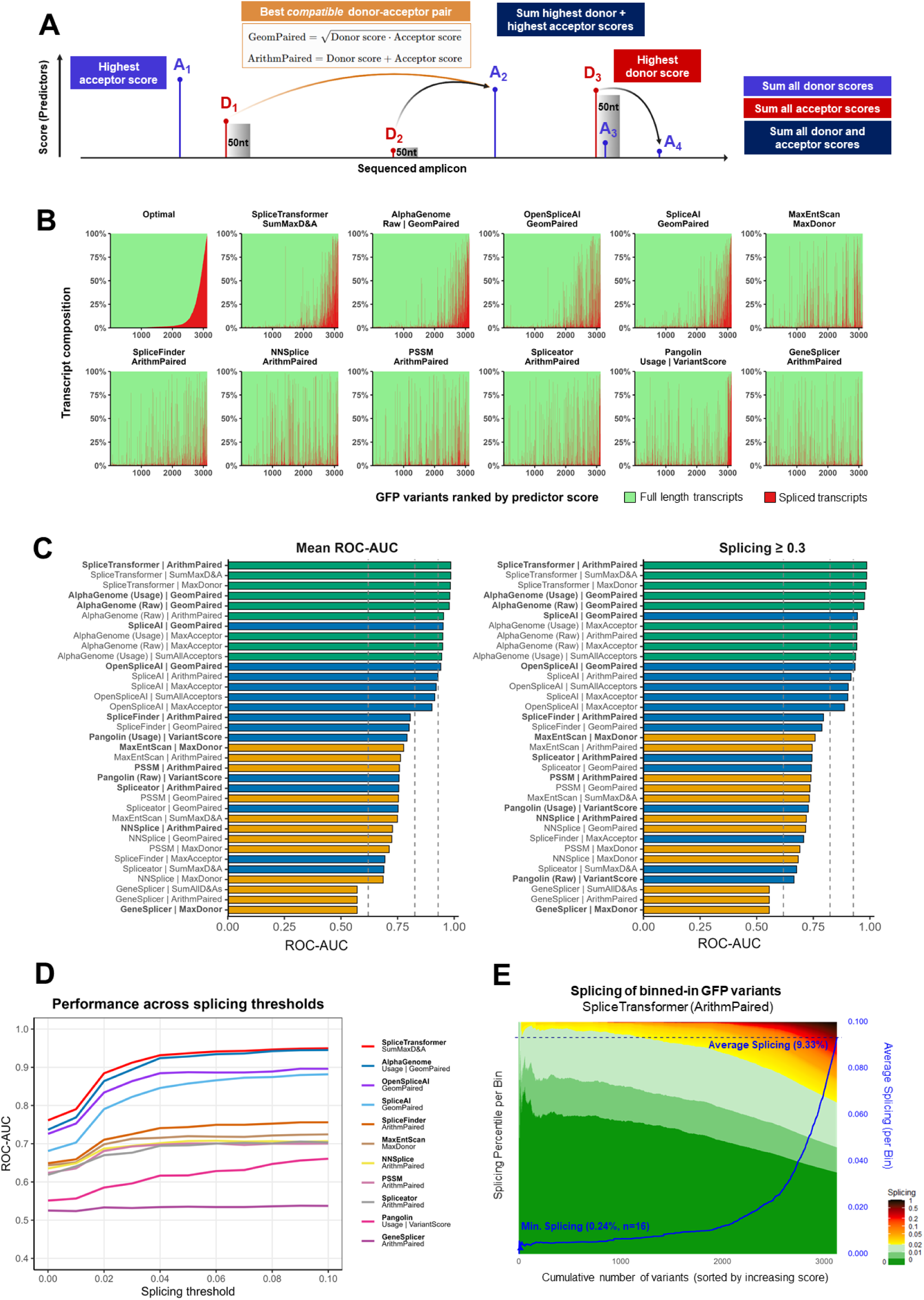
Gene-level prediction of splicing propensity. **(A)** Schematic of the eight gene-level scoring strategies evaluated. **(B)** Ranking of GFP variants by gene-level splicing propensity across prediction methods. Variants were ordered by their experimentally measured splicing (top left) or by predicted splicing, and transcript composition is shown for each gene as the fraction of full-length (green) and spliced (red) transcripts. For each predictor, only the highest-performing gene-level scoring strategy is shown, indicated above each panel. **(C)** Benchmarking of gene-level splicing prediction across scoring strategies: mean ROC-AUC across thresholds (left) and performance at high splicing activity (splicing ≥ 0.3; right). For clarity, only the three highest-performing scoring strategies per predictor are displayed. **(D)** Classification performance across the low-splicing regime (0–0.1 splicing thresholds). Curves show ROC-AUC across progressively increasing splicing thresholds; for each predictor, only the highest-performing gene-level scoring strategy is shown. **(E)** Gene-level prioritization landscape generated using SpliceTransformer ArithmPaired. GFP variants were ordered by increasing predicted splicing (X axis). The filled landscape (left Y axis) shows the distribution of experimentally measured splicing across variants up to and including the focal variant, coloured by splicing level (see scale); the blue curve (right Y axis) shows the cumulative average splicing across all variants up to and including the focal variant. The lowest average splicing among cumulative subsets of at least 10 variants was 0.24% (16 lowest-scoring variants), compared with a full-library average of 9.3%.

As an initial assessment, we examined the ability of each predictor’s best-performing strategy to rank GFP constructs by their splicing propensity. In Fig. 5B and Fig. S7, GFP variants are ranked either by their observed splicing propensity (top left) or by a given aggregated score, with vertical columns representing the fraction of spliced (red) and unspliced (green) reads per construct. Performance varied substantially between predictors, with SpliceTransformer and AlphaGenome showing the strongest agreement with the empirical ordering.

We next evaluated the ability of each predictor to classify constructs as spliced or non-spliced across a range of splicing thresholds, focusing on three representative values: 0%, 10%, and 30% (Fig. 5C, Suppl. Fig. 8). SpliceTransformer achieved the highest overall performance (average AUC 0.89), followed closely by AlphaGenome (0.88), the other deep learning models (0.83–0.84), motif-based and lightweight predictors (0.68–0.73), and GeneSplicer (0.53). SpliceTransformer was also the predictor least sensitive to aggregation strategy, whereas the other models strongly depended on aggregation strategy for their performance (Suppl. Fig. 9). No single aggregation strategy was best for all predictors: Pairing-based scores gave the highest AUC for most predictors, with GeomPaired working best with most deep-learning models (SpliceAI, OpenSpliceAI, AlphaGenome) and ArithmPaired for motif-based and lightweight models (PSSM, NNSplice, Spliceator, SpliceFinder). This indicates that most models benefit from the explicit donor–acceptor pairing that a single-site score does not capture. For a minority of predictors, a per-site maximum was sufficient: MaxEntScan and GeneSplicer performed best using the maximum donor score alone.

Performance was generally higher at splicing thresholds above 0.02 (Fig. 5D). This may reflect a limitation of the experimental ground truth: variants with no observed splicing could include false negatives arising from our coverage threshold of 50 reads per variant, which would reduce apparent performance at the lowest threshold. Overall, deep-learning models outperformed motif-based models in the whole-gene prediction benchmark, and the best aggregation strategy was predictor-dependent rather than universal.

### Implications for splicing-aware construct design

To explore the applicability of splice predictors in a gene engineering context, we simulated a scenario in which a collection of candidate variants is computationally screened to identify those with a low splicing propensity. After ranking GFP variants by their SpliceTransformer ArithmPaired score, we represented the experimentally observed splicing outcome of each variant with a colour scale, from dark green (unspliced) to dark red and black (fully spliced) (Fig. 5E). This revealed effective prioritisation of variants with null or low observed splicing. However, some level of splicing (1–2% per gene) was observed even among lowest-scoring variants, suggesting that computational screening might not be sufficient to fully eliminate splicing among randomly recoded genes. We further evaluated a scenario in which a set of 25 GFP lowest-scoring variants from each predictor are selected for experimental validation. Sets of variants identified by SpliceTransformer and AlphaGenome consistently showed the lowest gene-level splicing, whereas other predictors included constructs with higher splicing levels among their top candidates (Suppl. Fig. 10).

Finally, we asked whether splicing predictors can be used to identify the specific sequence determinants underlying splicing risk and guide targeted sequence optimization. To evaluate this, we applied in silico saturation mutagenesis to a GFP variant that shows a high level of splicing. We first confirmed that SpliceTransformer accurately captures the observed donor and acceptor sites for the selected variant. Consistent with Nanopore measurements (Fig. 6), the model assigned the highest donor and acceptor scores to the splice sites responsible for the vast majority of splicing events. For each position, we then generated all possible single-nucleotide substitutions, and each mutated sequence was scored using SpliceTransformer to quantify changes in predicted splice-site strength. Across the resulting mutational landscape, substitutions produced a broad spectrum of effects, ranging from minimal impact to substantial reductions in predicted splice-site strength (Fig. 6). Most substitutions did not generate stronger alternative splice sites than the observed configuration. However, a subset of mutations altered splice-site predictions, either weakening dominant sites or enabling the emergence of alternative splice signals. Notably, even substitutions at positions distal from the observed splice sites were capable of reconfiguring splice-site usage and promoting alternative splicing signals. These results suggest that computational predictors can identify minimally spliced variants and prioritise sites for targeted mutagenesis in gene engineering.

**Figure 6.**
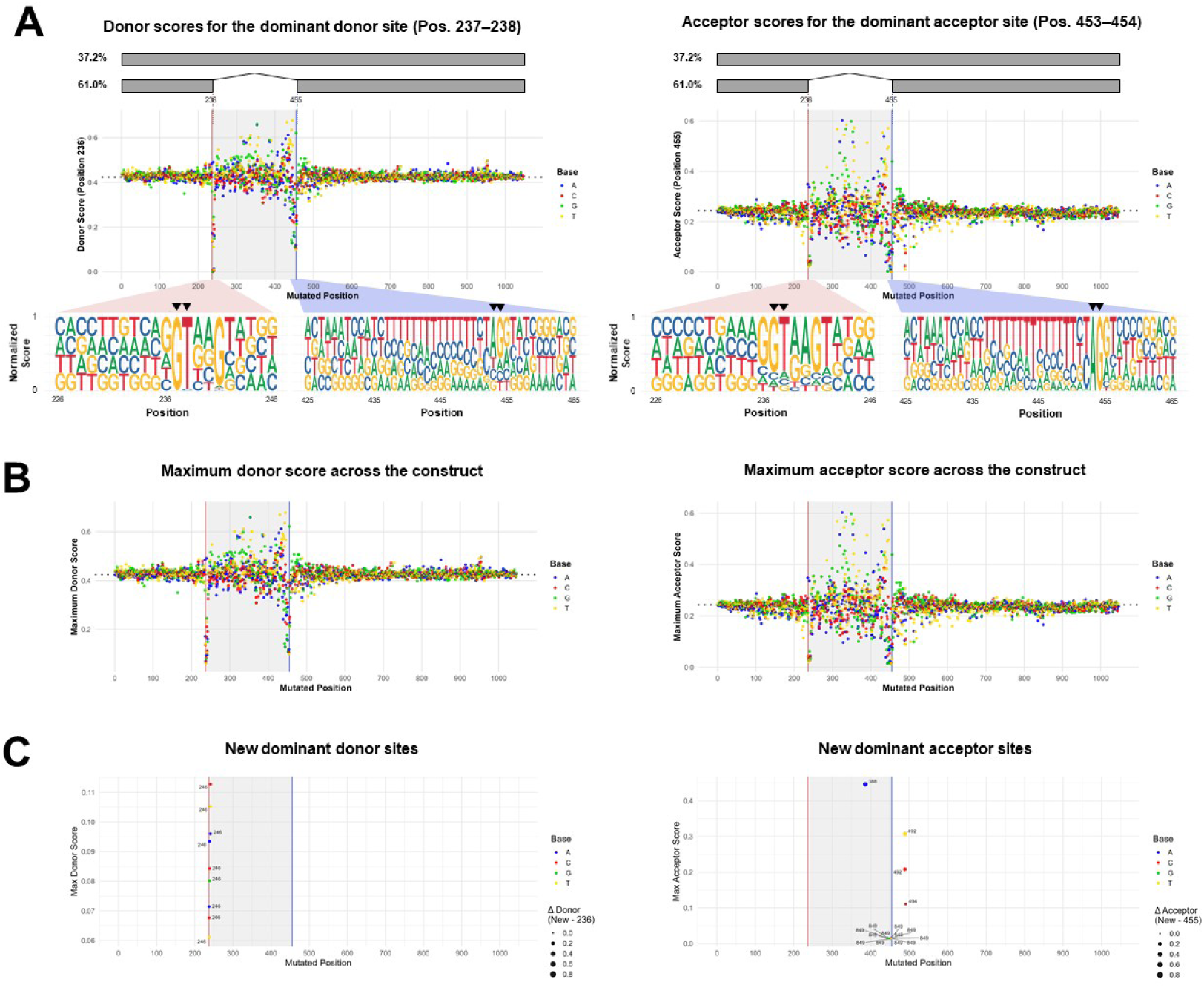
SpliceTransformer-guided saturation mutagenesis reveals sequence determinants of dominant cryptic splice sites. (A) In silico saturation mutagenesis of a representative GFP construct. Upper bars show the two most abundant transcript isoforms detected by Nanopore sequencing with their relative frequencies. Black arrowheads indicate the dominant donor and acceptor splice sites analysed below. For each position, all possible single-nucleotide substitutions were generated and scored using SpliceTransformer. Scatter plots show the predicted scores for the dominant donor (positions 237–238) and acceptor (positions 453–454) sites following each substitution; coloured dots indicate the substituted nucleotide, and dashed horizontal lines denote the score of the unmutated sequence. Sequence logos summarize the contribution of individual nucleotides to splice-site strength within the local context, with logo heights derived from normalized mutation-dependent scores, such that nucleotides whose substitution most strongly reduced predicted splice-site strength contribute proportionally greater information content. (B) Maximum predicted donor and acceptor scores anywhere within the construct following each single-nucleotide substitution. (C) Mutations that changed the identity of the dominant donor or acceptor site within the construct. Only substitutions shifting the dominant site are shown, with labels indicating the positions of the newly dominant sites. Point size reflects the score difference between the newly dominant and the original dominant donor (positions 237–238) or acceptor (positions 453–454) site.

## Discussion

Here, we ranked eleven splicing predictors across thousands of diverse variants of a synthetic reporter gene, using two types of benchmarks: position-level prediction of splice site usage and construct-level splicing propensity. We find that the ranking is largely stable across benchmarks — SpliceTransformer ranks highest, closely followed by AlphaGenome. SpliceAI and OpenSpliceAI also perform well, while tools that ignore long-range context show moderate performance. This largely reflects low specificity: these methods assign high scores to many positions that are not used as splice sites, so although they recover genuine sites, they also report many false positives. These findings agree with earlier benchmarking efforts using massively parallel assays and clinical variants. Smith and Kitzman benchmarked eight predictors against massively parallel splicing assays in five genes, finding deep-learning tools best overall and SpliceAI and Pangolin most sensitive [20]. More recently, large-scale saturation mutagenesis of more than half a million variants across hundreds of human exons similarly demonstrated strong performance of modern deep-learning predictors, with SpliceAI and Pangolin emerging as the top-performing models in that evaluation setting [37]. Similarly, clinical benchmarking studies identified SpliceAI as the strongest individual predictor for prioritising splice-disrupting variants and demonstrated that integrating splicing predictors into diagnostic workflows can substantially improve diagnostic yield [38]. A benchmark of pretrained genomic language models ranked SpliceTransformer among the strongest splice-site predictors, while noting that any model’s advantage depends on context length, training data, and task [39]. Nevertheless, our ranking also diverges from some earlier reports. AlphaGenome was outperformed by Pangolin on a minigene reporter assay in its own benchmarking [4], yet ranks clearly ahead of Pangolin on our dataset — indicating that relative standing can shift between evaluation settings. SpliceTransformer, meanwhile, was absent from many earlier comparisons, leaving its standing unclear; our benchmark places it first across tasks.

Beyond ranking predictors, our data allow their scores to be interpreted in terms of splicing outcomes. A raw predictor score is not in itself an expected level of splicing, and relating the two has so far been done one tool at a time and in each tool’s own units. Pangolin produces a quantitative estimate of site usage [15]; OpenSpliceAI calibrates its output probabilities against empirical ones [17]; and dedicated quantitative-usage models report that SpliceAI tends to overestimate usage [40]. Each of these calibrations is specific to a single predictor, so scores are not comparable across tools: a SpliceAI score of 0.2 and a SpliceTransformer score of 0.2 do not correspond to the same splicing outcome, and the SpliceAI delta-score thresholds in common clinical use (0.2, 0.5, 0.8) [14] do not transfer to other methods. We address this by mapping every predictor’s raw scores onto a single empirical axis, the measured fraction of spliced reads at each position, using generalised additive models (Fig. 4). This provides a unified calibration in which a given splicing outcome corresponds to a defined score in each tool.

High performance in our benchmark does not necessarily mean that a tool is best-suited for all applications. For example, splice predictors are commonly used for the identification of pathogenic variants [38, 41, 42], with variants predicted to change splicing relative to the reference sequence deemed to be pathogenic. In this setting, splicing predictors can use additional signals that are not present in our synthetic dataset but are relevant for pathogenicity: for example, evolutionary conservation at the single nucleotide level, sequence variation across the population, or patterns of splicing observed across human tissues. A tool that has been trained to take advantage of these signals can perform relatively better at pathogenicity prediction than in our synthetic benchmark. Recent work has suggested that strong performance on held-out genomic sequences does not necessarily imply equivalent performance on entirely novel sequence contexts, because related sequences may still be shared between training and evaluation datasets [43]. Although motif-based and early machine-learning methods showed only moderate performance in our study, they remain valuable in applications focused on the analysis of individual splice regulatory elements. For example, when the goal is to evaluate the intrinsic strength of a specific splice donor or acceptor motif, rather than to predict transcript-level splicing outcomes, local motif-based models may be more informative than context-aware deep learning approaches, as their scores directly reflect the properties of the motif being examined rather than integrating information across a broader sequence context.

We evaluated all predictors as released, without fine-tuning them on our data. Although fine-tuning can substantially improve performance on a given task [39], our aim is to test how models generalise to unseen, out-of-distribution sequences as they are actually deployed, not after adaptation to our data. For the same reason, we used each tool with its default settings, which may partly explain the near-chance performance of GeneSplicer; tuning model parameters could improve performance, but doing so selectively would not reflect typical use.

Our synthetic benchmark is particularly relevant for gene engineering applications in which different fragments of DNA are fused, codon optimised, and otherwise altered, potentially generating splice regulatory signals that may lead to cryptic splicing. Recent work indicates that such events are more common than previously thought: cryptic splicing has been documented across diverse plasmid-expressed transgenes despite the absence of annotated introns [44], can be driven systematically by defined sequence features such as U-rich elements [45], and, as we recently showed, affects a large fraction of heavily recoded synonymous constructs [35]. The consequences for expression can be substantial. For example, the SARS-CoV-2 Spike ORF contains numerous predicted cryptic splice sites and undergoes extensive aberrant splicing when expressed from DNA vectors in the nucleus, generating truncated splice isoforms that lack the transmembrane domain and may be secreted. Furthermore, codon optimization can increase the abundance of predicted splice sites and splicing events [33]. Splicing predictors have begun to be used to identify and eliminate unwanted splice sites during sequence engineering, and have recently been incorporated into generative frameworks that design entirely new synthetic sequences with programmed splicing behaviour and cell-type specificity [46]. Across these applications, the choice of predictor matters, and our benchmark indicates which architectures are most reliable when sequences depart from the natural genome: transformer-based and other context-aware deep-learning models substantially outperform motif-based tools, and are the better default for screening engineered constructs.

## Methods

### Synthetic GFP benchmarking datasets

Benchmarking analyses were performed using two previously published GFP libraries described by Ahmad et al. The first was a pooled library of 4,675 synonymous GFP variants (GFP_optimized) generated through large-scale third-codon randomization while preserving the encoded amino acid sequence. Degenerate codon selection was biased towards codons associated with elevated GC content and codon adaptation, producing variants representative of sequence design strategies commonly used in recombinant protein production, gene therapy, and synthetic biology. An independent validation dataset consisted of an arrayed design-of-experiments GFP library (GFP_DOE) comprising 194 variants generated through factorial combinations of GC content, CpG dinucleotide frequency, and codon adaptation index (CAI), thereby systematically exploring a broader sequence landscape.

Transcriptomic measurements for both libraries were from Ahmad et al. Briefly, long-read Oxford Nanopore sequencing experiments were performed on RNA isolated from transiently transfected HeLa cells. Raw sequencing data, transcript annotations, and splice junction calls were obtained from the original study and re-analysed for benchmarking purposes. Only variants supported by at least 50 UMI-corrected reads were retained for downstream analyses, resulting in 3,125 variants from the GFP_optimized library and 189 variants from the GFP_DOE library. Donor and acceptor splice-site usage were quantified at nucleotide resolution as the fraction of transcript molecules in which a given position participated in a splice junction relative to all transcript molecules assigned to the corresponding construct.

### Prediction methods

#### Position-specific scoring matrix (PSSM)

Donor and acceptor splice-site strengths were evaluated using custom position-specific scoring matrices (PSSMs) derived from ∼90,000 annotated human splice junctions using windows of [−3, +6] around GT donor sites and [−14, +1] around AG acceptor sites, respectively. All potential donor and acceptor sites within each GFP variant were scored using a sliding-window approach, retaining only canonical GT donor and AG acceptor motifs for downstream analyses. PSSM matrices are provided in the project GitHub repository.

### MaxEntScan

Splice-site strengths were evaluated using MaxEntScan, a maximum entropy-based splice-site prediction framework. Each GFP sequence was scanned using overlapping 9-nt donor windows and overlapping 23-nt acceptor windows, consistent with the sequence requirements of MaxEntScan. All possible windows for which the required motif length could be extracted were scored.

### NNSplice

Splice-site strengths were evaluated using NNSplice v0.9, a neural network-based splice-site prediction framework. Predictions were obtained using the Berkeley Drosophila Genome Project (BDGP) implementation with the “Human/other” model, scoring both donor and acceptor splice sites. Minimum donor and acceptor score thresholds were set to 0 to retain all candidate predictions, and reverse-strand prediction was disabled.

### GeneSplicer

Splice-site strengths were evaluated using GeneSplicer with the human splice-site model. Donor and acceptor predictions were generated using donor and acceptor score thresholds of 0 (-d 0, -a 0). Scores were reported only for positions classified as candidate donor or acceptor splice sites by GeneSplicer rather than for every nucleotide position in the sequence.

### SpliceFinder

SpliceFinder predictions were generated using the published convolutional neural network model. Each GFP sequence was scanned using overlapping 400-nt windows with a stride of 1 nt, consistent with the window size used during model training. For each window, the model returned donor-site, acceptor-site, and non-splice-site probabilities. Donor and acceptor probabilities were assigned to the central nucleotide position of each window, thereby generating nucleotide-resolution predictions across the sequence. Because predictions were assigned to the centre of a 400-nt window, only positions that could serve as the central nucleotide of a complete window were scored. Consequently, approximately the first and last 200 nt of each sequence were not assigned donor or acceptor scores.

### Spliceator

Spliceator predictions were generated using the published convolutional neural network model trained on multi-species splice junction datasets. Sequences were analysed using the default model parameters. Spliceator reports only positions classified as candidate donor or acceptor splice sites rather than generating scores for every nucleotide position in the input sequence. For each predicted splice site, the genomic position, splice-site type, and confidence score (0–1) were recorded. By default, only predictions with confidence scores greater than 0.5 are reported by the software.

### SpliceAI

Splice-site strengths were evaluated using the original SpliceAI implementation and its five pre-trained models. Each GFP sequence was padded with 5,000 Ns on both sides to provide the 10,000-nt sequence context required by the model. Predictions were generated by averaging outputs across the five SpliceAI models, and position-wise acceptor and donor probabilities were extracted for every nucleotide in each input sequence.

### Pangolin

Pangolin predictions were generated using a custom FASTA-based wrapper to obtain position-wise scores for each GFP variant. Sequences were padded with 5,000 Ns on both sides and analysed on the forward strand using the five-model ensemble for each Pangolin tissue-specific head. Both splice probability and usage outputs were computed for the four available tissues (heart, liver, brain, and testis) and averaged across tissues at each nucleotide position. Throughout the manuscript, the resulting tissue-averaged *splice probability scores* and *usage scores* are referred to as *Pangolin Raw* and *Pangolin Usage*, respectively. Because Pangolin does not provide separate donor and acceptor scores, Pangolin was excluded from donor- and acceptor-specific splice-site benchmarking analyses. Donor–acceptor pairing strategies could likewise not be applied. For Pangolin, construct-level scores were therefore defined as the maximum Raw score or maximum Usage score observed within each construct.

### SpliceTransformer

SpliceTransformer predictions were generated using the authors’ published implementation and pretrained model weights. Full-length coding sequences were provided in FASTA format and padded with 4,000 ambiguous nucleotides (N) on both sides prior to inference to provide the sequence context expected by the model. Following prediction, scores corresponding to the padding regions were discarded and only positions belonging to the original coding sequence were retained.

For each nucleotide position, the model produced splice-site prediction probabilities together with tissue-specific outputs. Based on the published output structure and empirical validation of the model output channels, per-position acceptor and donor probabilities were extracted from channels 1 and 2, respectively. No score thresholding was applied, and raw position-wise scores were retained for downstream analyses.

### OpenSpliceAI

OpenSpliceAI predictions were generated from custom GFP FASTA sequences using the pre-trained MANE 10000nt model (openspliceai-mane/10000nt). Predictions were run with a flanking-size parameter of 10,000 nt (-f 10000), corresponding to 5,000 nt of sequence context on each side, and no score filtering (-t 0.0), thereby retaining all donor and acceptor predictions for downstream benchmarking analyses.

### AlphaGenome

Splice-site strengths were evaluated using the AlphaGenome DNA model. GFP sequences were analysed using a 2,048-nt context window and were symmetrically padded with Ns to the required input length prior to prediction. For each nucleotide position, donor and acceptor splice-site probabilities were extracted from the SPLICE_SITES output, whereas donor and acceptor splice-site usage scores were extracted from the SPLICE_SITE_USAGE output. When multiple donor or acceptor tracks were available, the maximum value across the corresponding tracks was retained for each position. Both raw splice-site scores and splice-site usage scores were used for downstream benchmarking analyses.

### Benchmarking and calibration of splice-site prediction scores

#### Score harmonization and splice-site matching

To enable a fair comparison across splice-site prediction methods, scores were initially generated for all positions that could be evaluated by each predictor according to its respective input requirements. Because different tools use distinct coordinate conventions when reporting splice-site positions, method-specific coordinate offsets were applied where necessary to ensure that all scores referred to the same biological splice-site position. Unless otherwise stated, all downstream analyses were performed using harmonized coordinates.

Benchmarking analyses were restricted to canonical GT donor sites and AG acceptor sites. This focused performance evaluation on biologically relevant candidate splice sites and avoided classification results being driven primarily by the abundance of positions that are intrinsically incompatible with splicing. Positions lacking the corresponding motif were excluded prior to downstream correlation, ROC-AUC and precision–recall analyses.

### Benchmarking of splice-site prediction performance

Receiver operating characteristic (ROC) and precision–recall (PR) analyses were performed separately for donor and acceptor sites. Positions with experimentally measured splice-site usage exceeding predefined thresholds (0.02, 0.05, 0.10 or 0.50) were considered positive, whereas all remaining positions were considered negative. ROC-AUC and PR-AUC values were calculated using the pROC and PRROC R packages.

To evaluate performance across the full spectrum of splice-site activities, PR-AUC was additionally calculated across splice-site usage thresholds ranging from 0 to 1 in increments of 0.01. For each threshold, positions with experimentally measured splice-site usage exceeding the threshold were considered positive. Mean PR-AUC across all valid thresholds was used as a summary measure of overall predictor performance.

### Score calibration using generalized additive models

For score calibration, generalized additive models (GAMs) were fitted separately for each predictor and splice-site type. Analyses were restricted to splice-compatible positions with non-missing experimental splice-site usage and non-missing predictor scores. Predictor scores were transformed prior to fitting: scores bounded at zero were log10-transformed after adding a pseudocount of 10^-3, whereas predictors with negative values were standardized.

For each predictor, separate binomial GAMs were fitted to model the probability that experimentally measured splice-site usage exceeded 0, 0.02, 0.10 or 0.30 as a smooth function of predictor score, using mgcv with a spline term s(score, k = 10). Predicted probabilities were obtained across 200 evenly spaced values spanning the observed score range. Category probabilities were then derived from the cumulative threshold probabilities, corresponding to no detectable splicing, 0–0.02, 0.02–0.10, 0.10–0.30 and >0.30 splice-site usage.

### Construct-level prediction strategies and benchmarking

#### Construct-level score aggregation strategies

To derive a single splicing propensity score per gene, position-level donor and acceptor predictions were aggregated using multiple gene-level scoring strategies. The maximum donor score (MaxDonor) and maximum acceptor score (MaxAcceptor) corresponded to the highest-scoring donor or acceptor site within each gene, respectively. Additional strategies included the sum of all donor scores (SumAllDonors), the sum of all acceptor scores (SumAllAcceptors), the sum of all donor and acceptor scores (SumAllD&As), and the sum of the maximum donor and maximum acceptor scores (SumMaxD&A).

Donor–acceptor pairing strategies were designed to approximate productive splice junction formation. For each construct, the five highest-scoring donor sites were considered as candidate splice donors. For each donor, the highest-scoring acceptor located at least 50 nucleotides downstream was selected. The 50-nt minimum distance was introduced to avoid biologically implausible donor–acceptor pairings and to approximate the spacing generally required for productive splicing. Donor and acceptor scores were then combined either using their geometric mean (GeomPaired) or their arithmetic sum (ArithmPaired). The highest-scoring donor–acceptor pair was retained as the gene-level score.

### Benchmarking of gene-level prediction performance

Gene-level prediction performance was evaluated by comparing experimentally measured total splicing per GFP variant with each gene-level score. ROC-AUC analyses were performed for each predictor–strategy combination. Genes with experimentally measured splicing greater than 0, 0.10 or 0.30 were considered positives, whereas all remaining constructs were considered negatives. To summarize performance across the full range of gene-level splicing activities, ROC-AUC was additionally calculated across thresholds from 0 to 0.99 in increments of 0.01, and the mean across valid thresholds was reported.

For the low-splicing regime, ROC-AUC was calculated across thresholds from 0 to 0.10 in increments of 0.01. Missing, infinite or non-variable score sets were excluded from ROC-AUC calculations.

### In silico saturation mutagenesis

To investigate sequence determinants of cryptic splice-site recognition, in silico saturation mutagenesis was performed on a representative GFP construct exhibiting 62.8% splicing and a highly dominant splice isoform (>97% of all spliced transcripts). The dominant donor (GT positions 237–238) and acceptor (AG positions 453–454) splice sites identified by Nanopore sequencing were used as reference splice sites throughout the analysis.

Every possible single-nucleotide substitution was generated across the entire coding sequence by replacing each nucleotide with the three alternative bases. SpliceTransformer was then used to calculate donor and acceptor prediction scores across the entire sequence for each mutant.

Mutational effects were quantified by comparing mutant and wild-type scores at the dominant donor and acceptor sites, as well as by examining changes in the maximum donor and acceptor scores observed anywhere within the construct. Additional analyses identified mutations that generated alternative dominant donor or acceptor sites.

To summarize local sequence determinants of splice-site strength, mutation-derived scores were aggregated across all substitutions at each position and visualized as functional sequence logos. Nucleotide heights were derived from normalized mutation-dependent prediction scores, such that nucleotides whose substitution produced larger decreases in predicted splice-site strength contributed proportionally greater information content.

All statistical analyses and visualizations were performed in R (version 4.4.1) unless otherwise stated.

## Declarations

### Authors’ contributions

F.B.M. performed analyses, generated figures and wrote the manuscript. G.K. conceived the study, supervised the work and revised the manuscript. Both authors approved the final version.

## Supplementary Figures

**Supplementary Figure 1.**
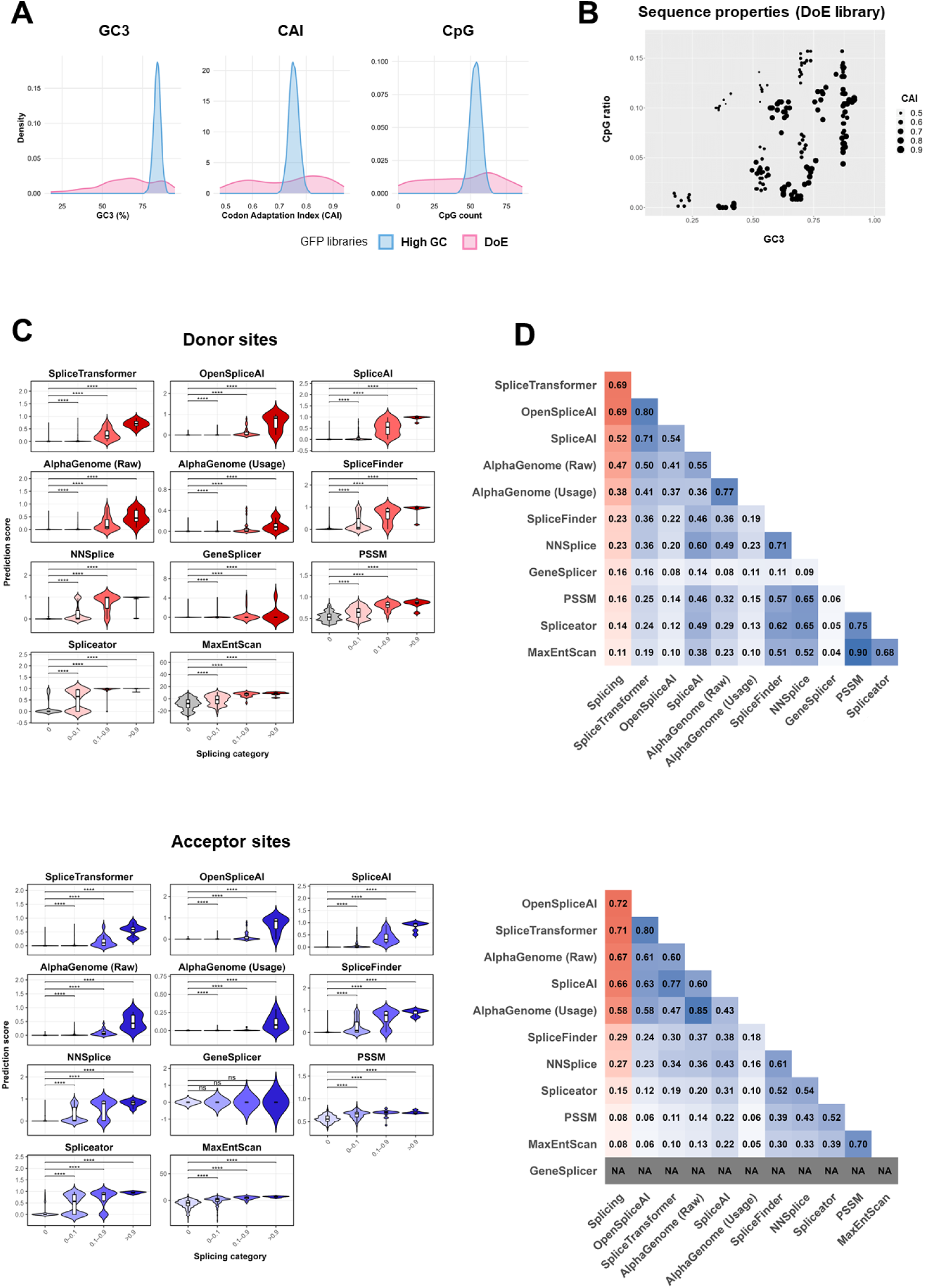
Generalization of splicing predictor performance across an independent GFP design-of-experiments (DoE) library. **(A)** Distribution of key sequence properties across the high-GC GFP library and the independent GFP DoE library, including GC3 content, codon adaptation index (CAI), and CpG dinucleotide frequency. **(B)** Joint distribution of GC3 content, CpG dinucleotide frequency, and codon adaptation index (CAI) across variants in the GFP DoE library, illustrating the broad exploration of sequence-property space enabled by the factorial design framework. **(C)** Distribution of predicted donor and acceptor splice-site scores across experimentally defined splicing activity bins in the GFP DoE library. **(D)** Pairwise Pearson correlation analysis between predicted splice-site scores and experimentally measured splice-site usage in the GFP DoE library (red column), as well as between prediction methods (blue tiles).

**Supplementary Figure 2.**
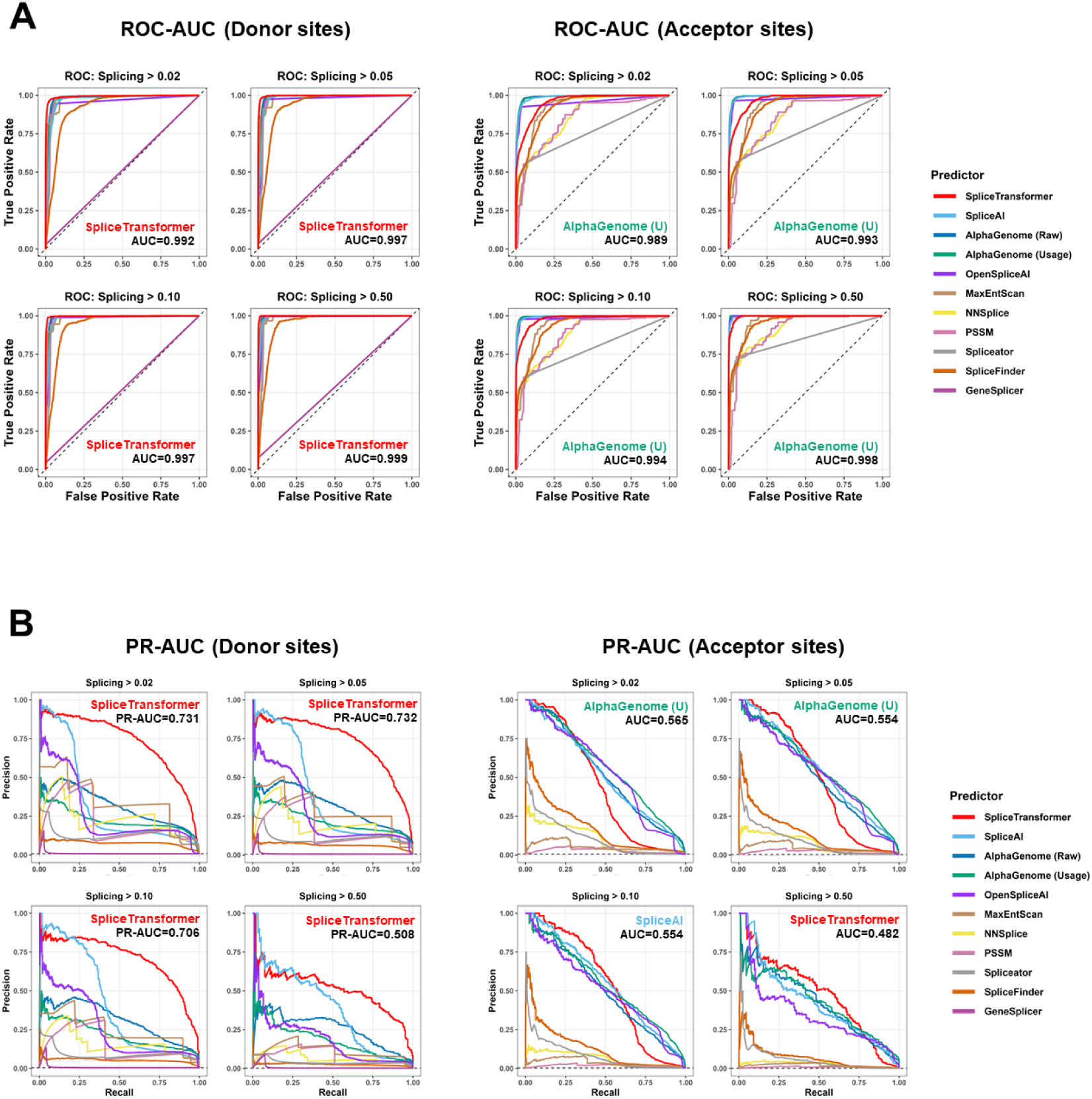
Extended ROC-AUC and precision–recall benchmarking across splice-site activity thresholds. **(A)** Receiver operating characteristic (ROC) curves for donor and acceptor splice-site classification across multiple experimentally defined splicing thresholds, ranging from low (>0.02) to highly active (>0.50) splice-site usage. **(B)** Precision–recall (PR) curves for donor and acceptor splice-site classification across experimentally defined splicing thresholds.

**Supplementary Figure 3.**
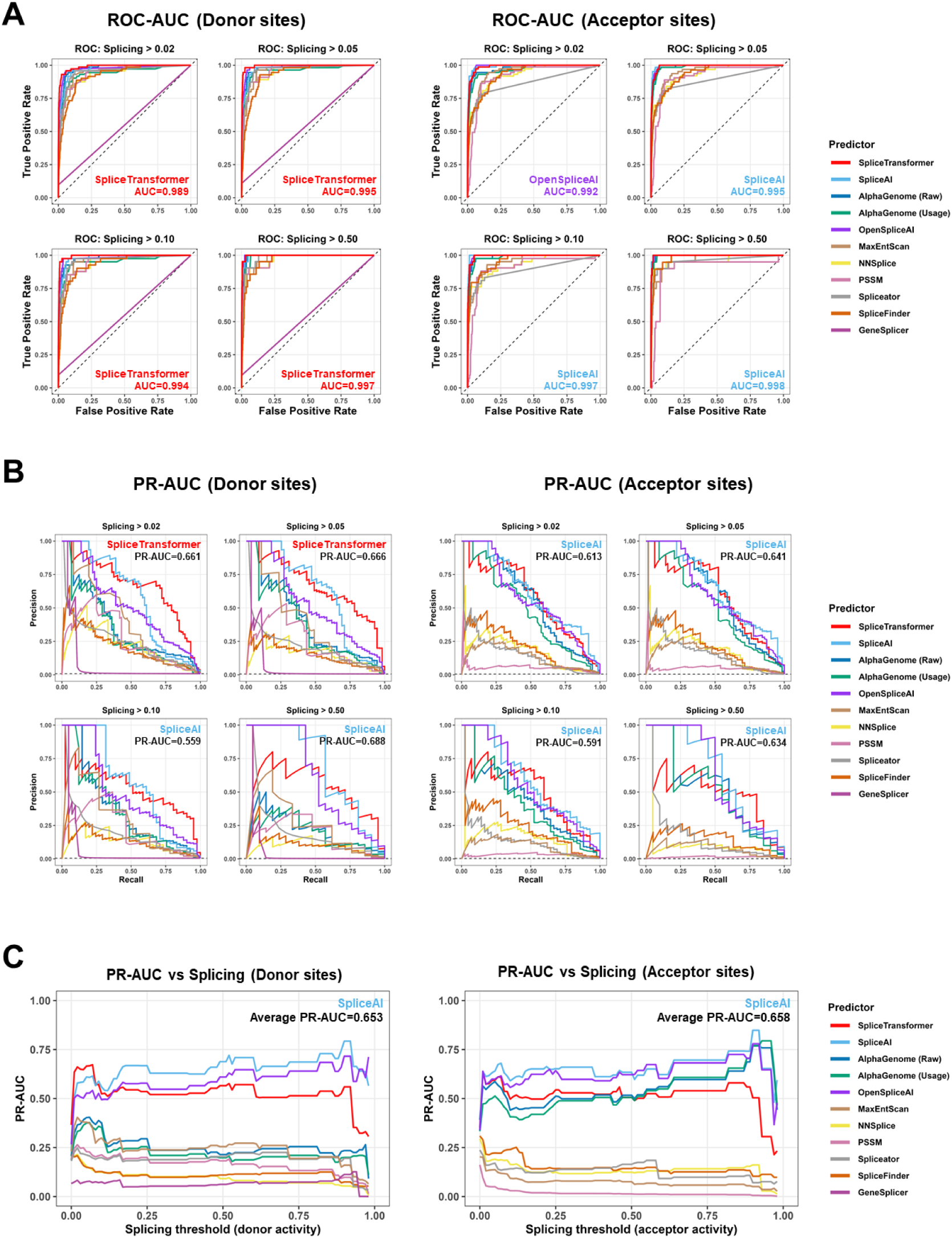
ROC-AUC and precision–recall benchmarking across splice-site activity thresholds in the independent GFP DoE library. **(A)** Receiver operating characteristic (ROC) curves for donor and acceptor splice-site classification across multiple experimentally defined splicing thresholds in the GFP design-of-experiments (DoE) library, ranging from low (>0.02) to highly active (>0.50) splice-site usage. **(B)** Precision–recall (PR) curves for donor and acceptor splice-site classification across the same range of experimentally defined splicing thresholds in the GFP DoE library. **(C)** PR-AUC values across a continuous range of experimentally measured splicing thresholds for donor and acceptor sites in the GFP DoE library.

**Supplementary Figure 4.**
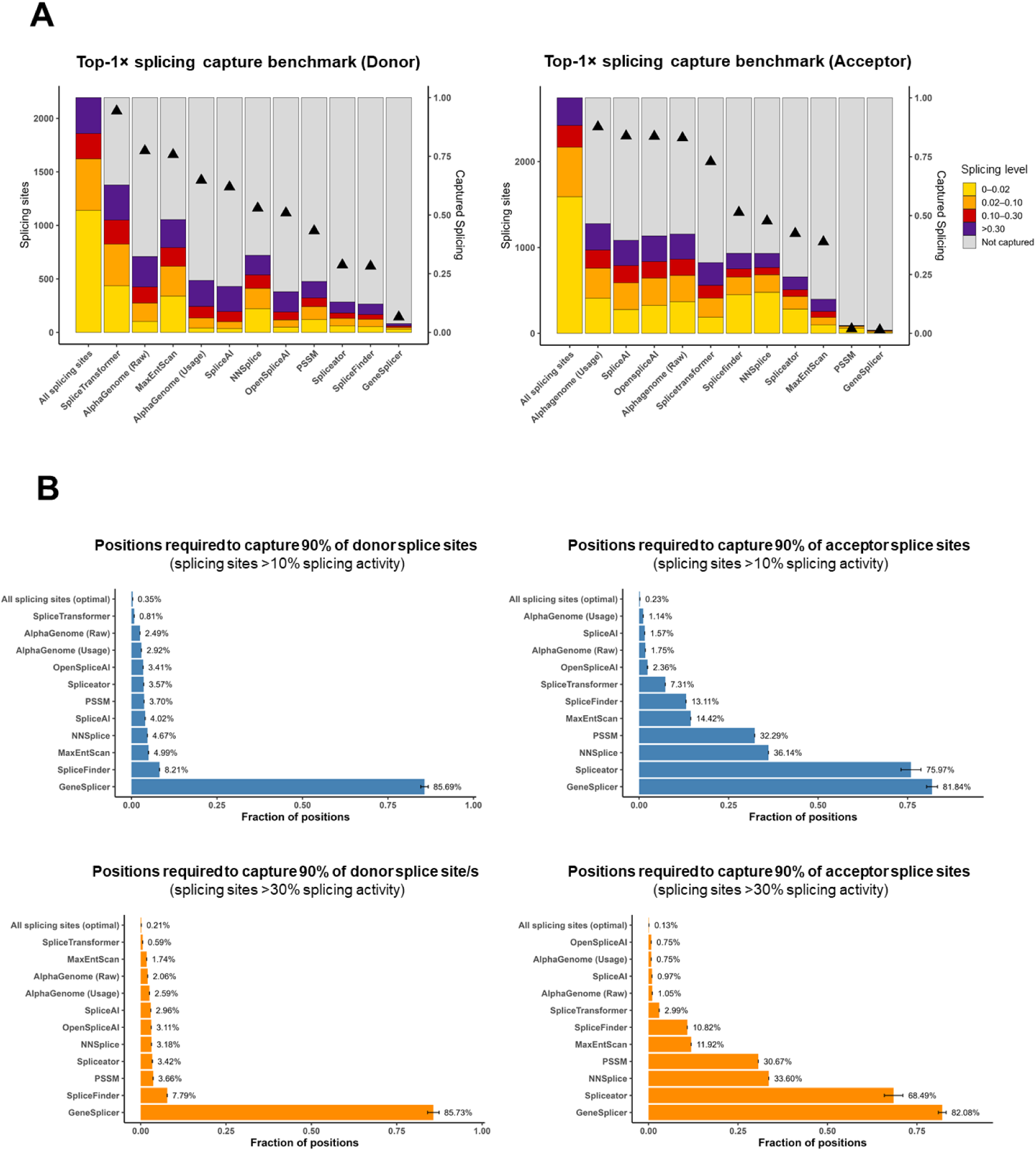
Deep learning predictors capture the majority of functional splice sites and enable calibration of splicing risk. **(A)** Constrained splice-site capture benchmarking for donor and acceptor prediction. For each predictor, splicing-compatible positions were ranked according to prediction score, and the top-scoring positions equal in number to the experimentally detected splice sites (*n*) were selected for benchmarking. Colored bars show the number of experimentally detected splice sites recovered within each splice-site activity class among the *n* top-ranked predicted positions, while black triangles indicate the cumulative fraction of total splicing activity recovered. Highly active splice sites (purple, red sections) were recovered at substantially higher rates than low or residual splicing events (yellow sections). **(B)** Bar plots show the fraction of splicing-compatible positions that must be prioritized to recover 90% of experimentally detected donor and acceptor splice sites with moderate-to-high (>10%) or highly active (>30%) splice-site usage. Theoretical optimal values (“All splicing sites”) correspond to the minimal fraction of positions represented by the experimentally detected splice sites themselves.

**Supplementary Figure 5.**
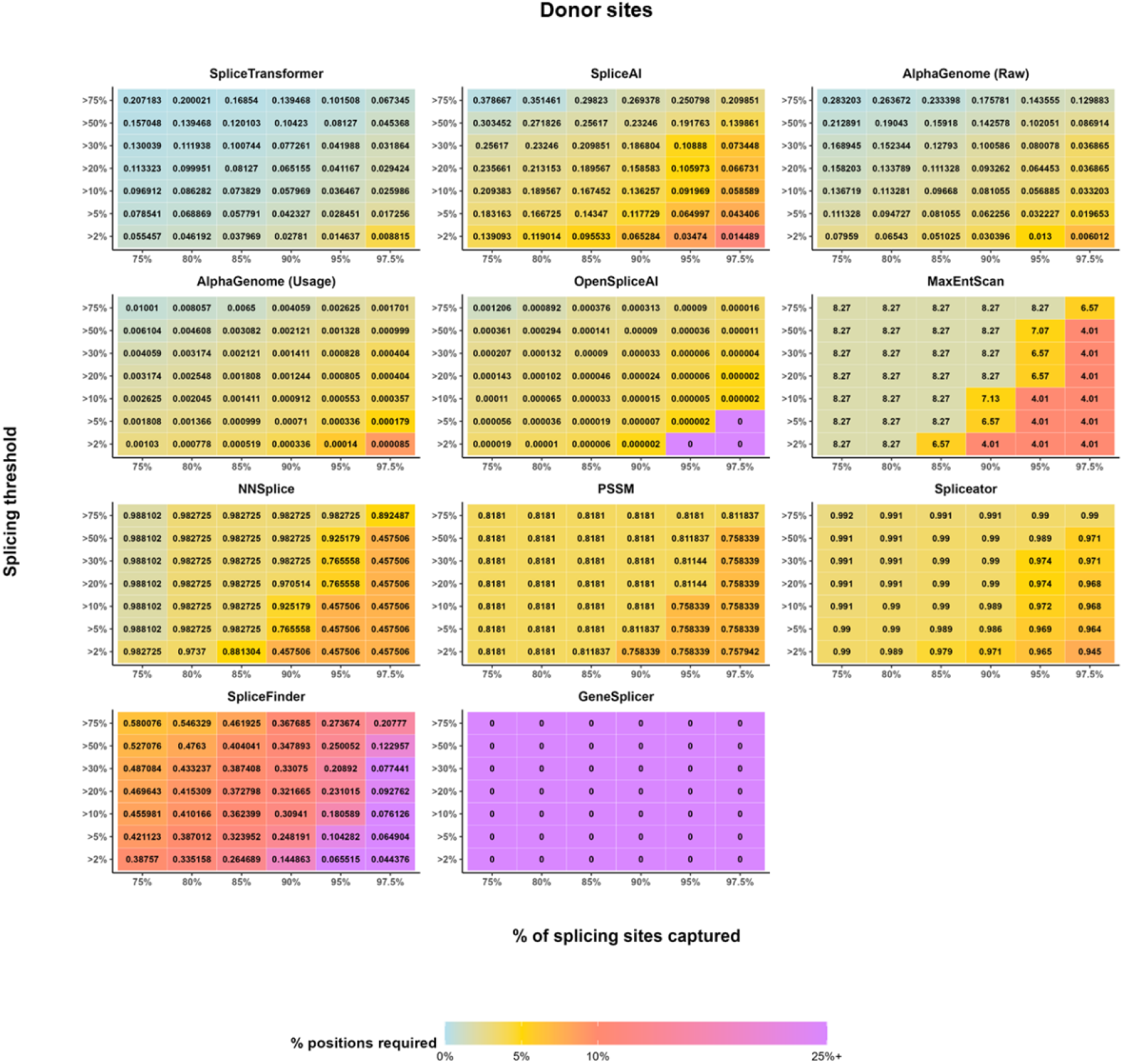
Score–coverage threshold matrices across all evaluated donor-site predictors. Heatmaps show the predictor score thresholds required to recover defined fractions of experimentally measured donor splice sites across different splice-site activity classes. The color scale represents the fraction of splicing-compatible positions that must be retained to achieve the corresponding coverage.

**Supplementary Figure 6.**
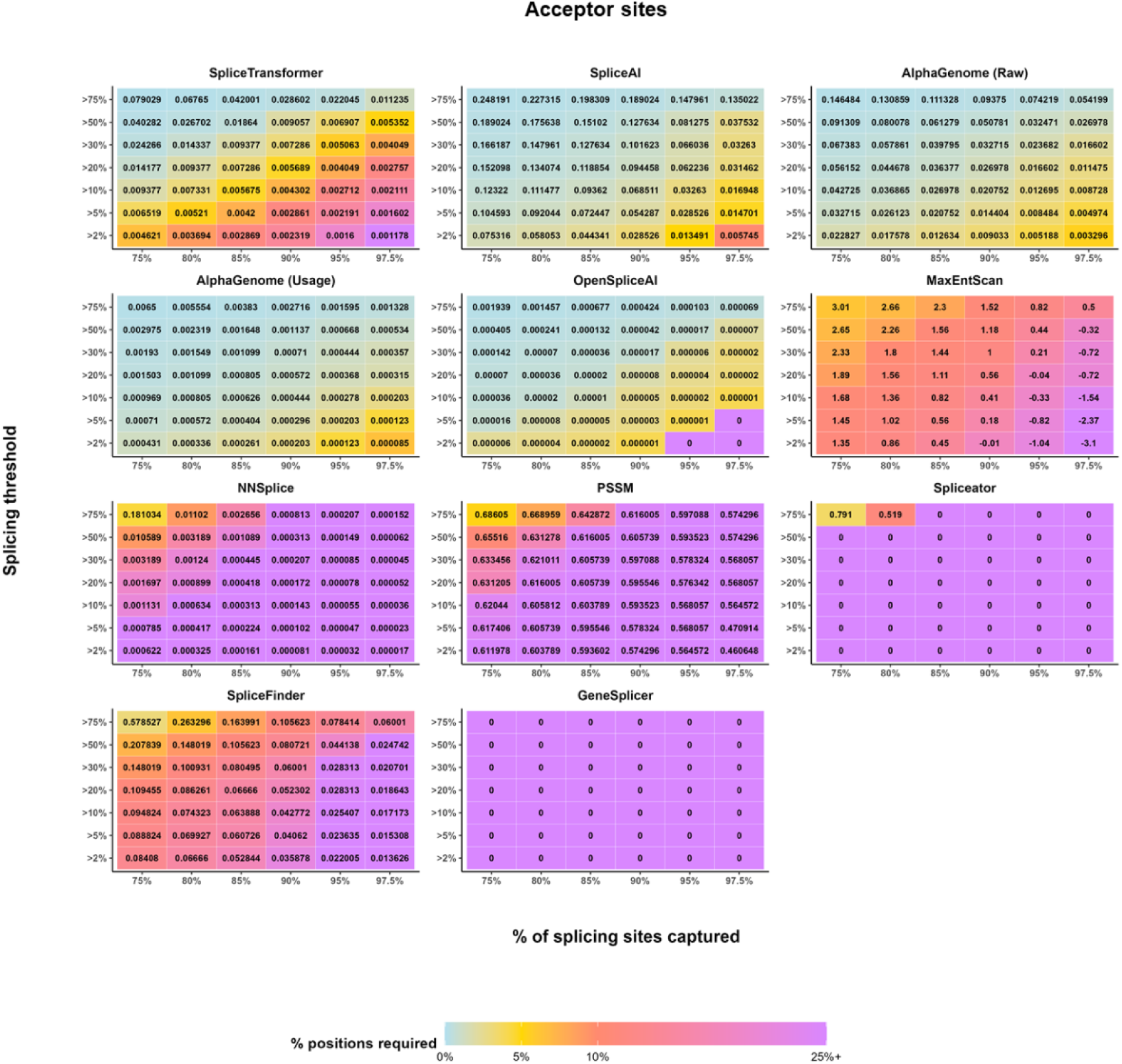
Score–coverage threshold matrices across all evaluated acceptor-site predictors. Heatmaps show the predictor score thresholds required to recover defined fractions of experimentally measured acceptor splice sites across different splice-site activity classes. The color scale represents the fraction of splicing-compatible positions that must be retained to achieve the corresponding coverage.

**Supplementary Figure 7.**
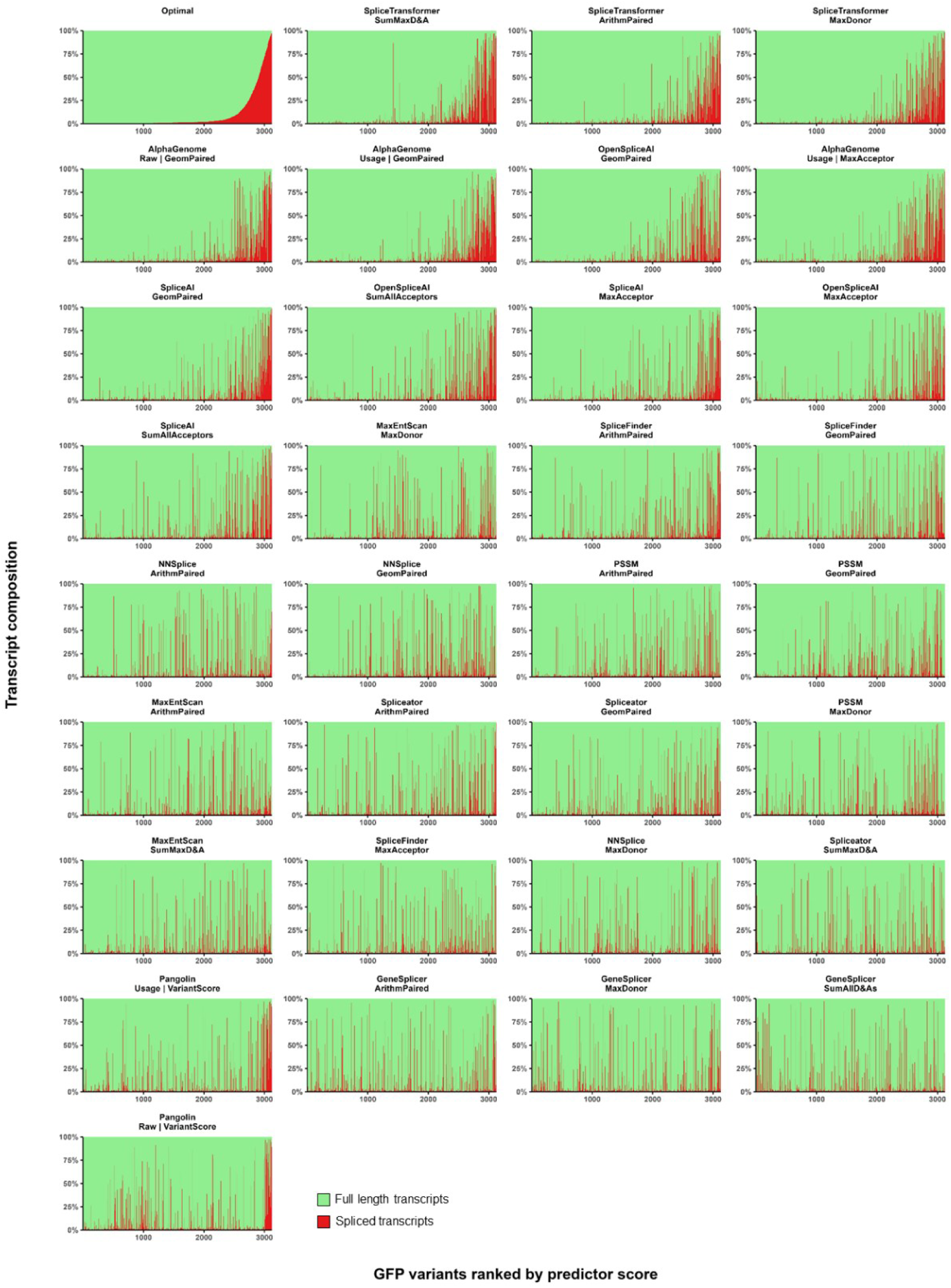
Gene-level ranking landscapes across evaluated prediction strategies. GFP variants were ordered according to increasing gene-level prediction score for each evaluated predictor and scoring strategy. The top-left panel ("Optimal") shows the empirical ordering obtained by ranking constructs according to their experimentally measured splicing activity and serves as a reference for ideal performance. For clarity, only the three scoring strategies with the highest Kendall’s τ-b values for each predictor are displayed. Predictors were ordered according to the performance of their best-scoring strategy, as measured by Kendall’s τ-b agreement between predicted gene-level scores and experimentally measured splicing activity. Vertical columns represent the fraction of full-length (green) and spliced (red) transcripts for each construct. Predictors with strong gene-level performance produced rankings that closely resembled the empirical ordering, with highly spliced constructs progressively accumulating toward the high-score end of the distribution.

**Supplementary Figure 8.**
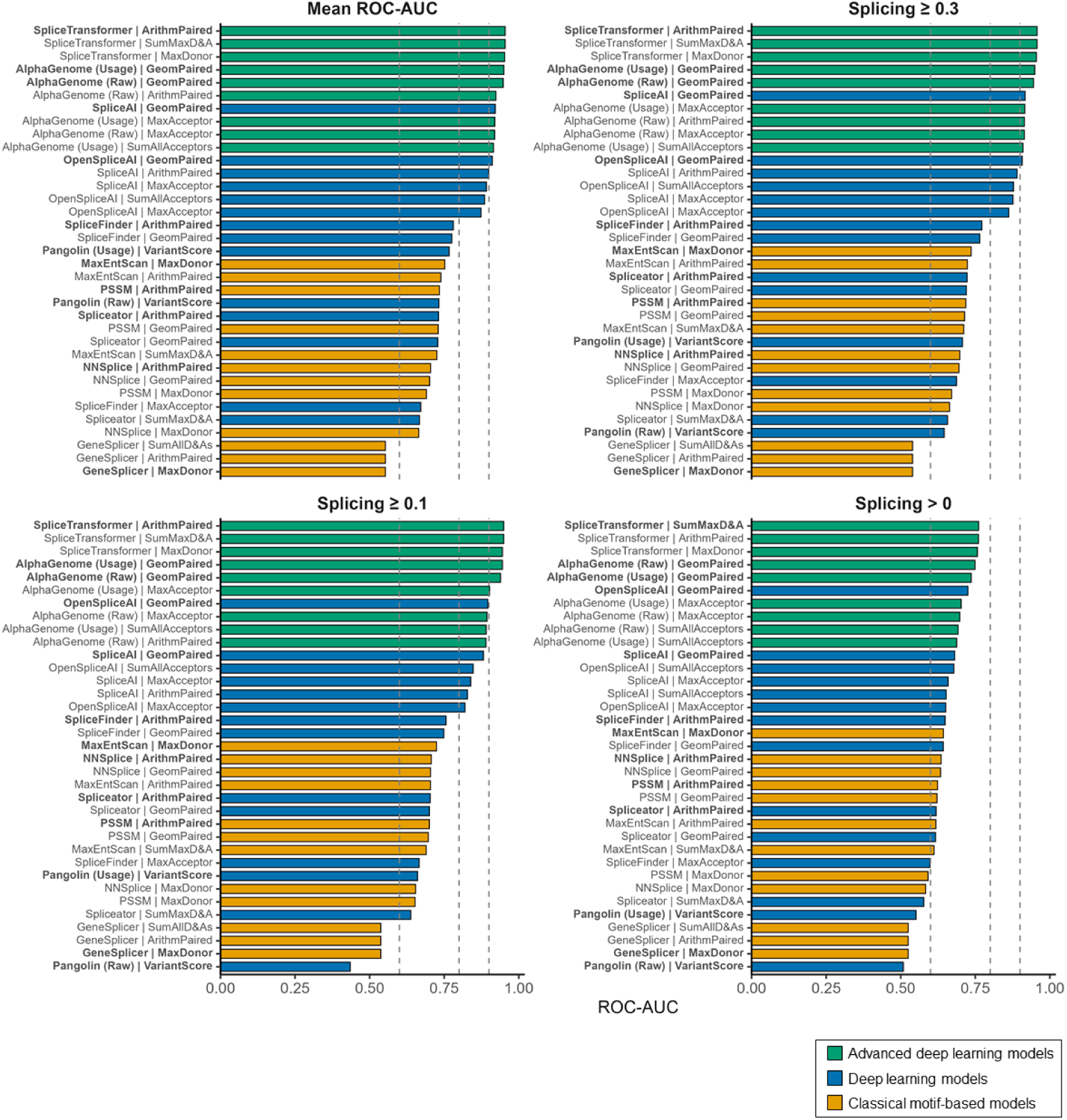
Extended gene-level ROC-AUC benchmarking across prediction strategies and splicing thresholds. Gene-level prediction strategies were ranked according to ROC-AUC performance across multiple splicing thresholds (0, 0.1, 0.3 and mean ROC-AUC across thresholds). For clarity, only the three highest-performing scoring strategies for each predictor are displayed. Predictors are grouped according to methodological class.

**Supplementary Figure 9.**
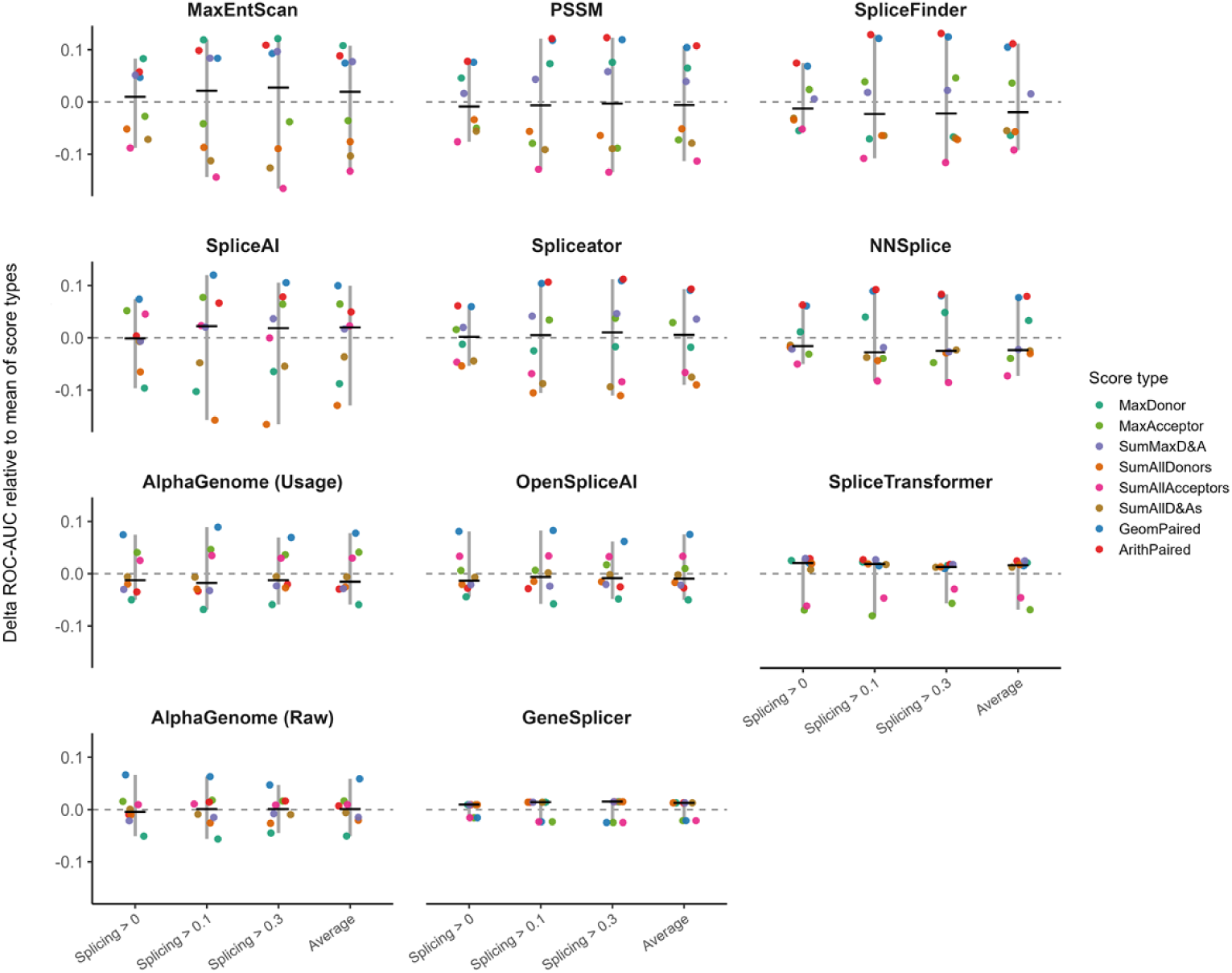
Effect of score aggregation strategy on predictor performance. For each predictor, residuals are shown relative to the mean ROC-AUC across all strategies for the same predictor and splicing threshold. Each colored point represents one aggregation strategy, the black bar indicates the median residual, and the grey vertical line spans the minimum and maximum residual values observed across strategies. Larger ranges indicate that predictor performance is more dependent on the choice of score aggregation strategy, whereas shorter ranges indicate greater robustness to the aggregation method. Positive residuals denote above-average performance and negative residuals denote below-average performance

**Supplementary Figure 10.**
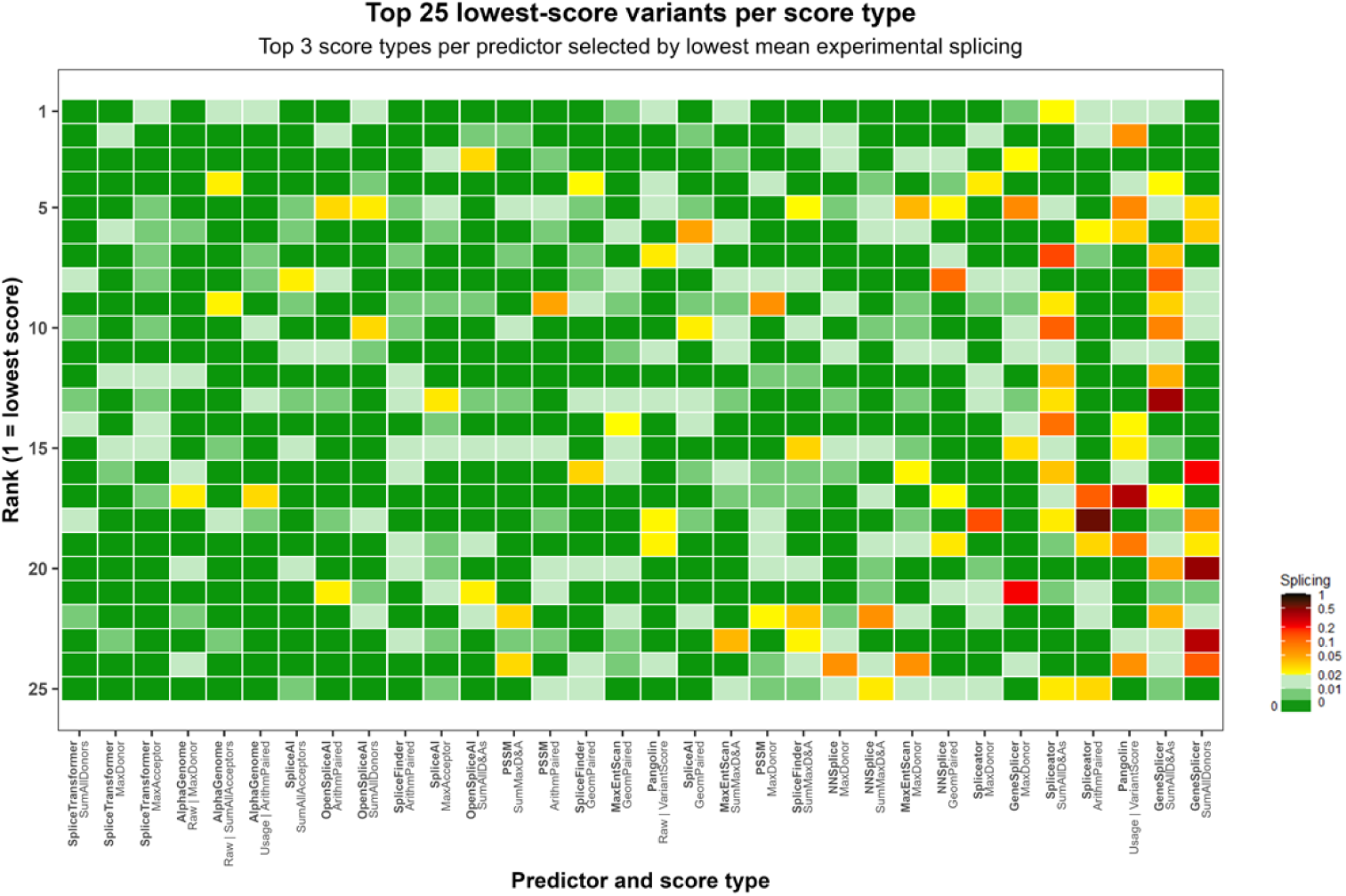
Splicing profiles of the lowest-scoring GFP variants across prediction methods. Heatmap showing experimentally measured splicing levels for the 25 lowest-scoring GFP variants identified by each predictor and scoring strategy. For clarity, for each predictor, only the three scoring strategies that produced the lowest average splicing among the selected 25 variants are displayed. Predictor–strategy combinations are ranked from left to right according to the average experimentally measured splicing of the selected variants.

## References

1. Jumper J, Evans R, Pritzel A, Green T, Figurnov M, Ronneberger O, Tunyasuvunakool K, Bates R, Zidek A, Potapenko A, et al: Highly accurate protein structure prediction with AlphaFold. Nature 2021, 596:583–589.

2. Vaishnav ED, de Boer CG, Molinet J, Yassour M, Fan L, Adiconis X, Thompson DA, Levin JZ, Cubillos FA, Regev A: The evolution, evolvability and engineering of gene regulatory DNA. Nature 2022, 603:455–463.

3. Cheng J, Novati G, Pan J, Bycroft C, Zemgulyte A, Applebaum T, Pritzel A, Wong LH, Zielinski M, Sargeant T, et al: Accurate proteome-wide missense variant effect prediction with AlphaMissense. Science 2023, 381:eadg7492.

4. Avsec Z, Latysheva N, Cheng J, Novati G, Taylor KR, Ward T, Bycroft C, Nicolaisen L, Arvaniti E, Pan J, et al: Advancing regulatory variant effect prediction with AlphaGenome. Nature 2026, 649:1206–1218.

5. Lee Y, Rio DC: Mechanisms and Regulation of Alternative Pre-mRNA Splicing. Annu Rev Biochem 2015, 84:291–323.

6. Matera AG, Wang Z: A day in the life of the spliceosome. Nat Rev Mol Cell Biol 2014, 15:108–121.

7. Rhine CL, Neil C, Wang J, Maguire S, Buerer L, Salomon M, Meremikwu IC, Kim J, Strande NT, Fairbrother WG: Massively parallel reporter assays discover de novo exonic splicing mutants in paralogs of Autism genes. PLoS Genet 2022, 18:e1009884.

8. Stormo GD, Schneider TD, Gold L, Ehrenfeucht A: Use of the ’Perceptron’ algorithm to distinguish translational initiation sites in E. coli. Nucleic Acids Res 1982, 10:2997–3011.

9. Yeo G, Burge CB: Maximum entropy modeling of short sequence motifs with applications to RNA splicing signals. J Comput Biol 2004, 11:377–394.

10. Reese MG, Eeckman FH, Kulp D, Haussler D: Improved splice site detection in Genie. J Comput Biol 1997, 4:311–323.

11. Pertea M, Lin X, Salzberg SL: GeneSplicer: a new computational method for splice site prediction. Nucleic Acids Res 2001, 29:1185–1190.

12. Wang R, Wang Z, Wang J, Li S: SpliceFinder: ab initio prediction of splice sites using convolutional neural network. BMC Bioinformatics 2019, 20:652.

13. Scalzitti N, Kress A, Orhand R, Weber T, Moulinier L, Jeannin-Girardon A, Collet P, Poch O, Thompson JD: Spliceator: multi-species splice site prediction using convolutional neural networks. BMC Bioinformatics 2021, 22:561.

14. Jaganathan K, Kyriazopoulou Panagiotopoulou S, McRae JF, Darbandi SF, Knowles D, Li YI, Kosmicki JA, Arbelaez J, Cui W, Schwartz GB, et al: Predicting Splicing from Primary Sequence with Deep Learning. Cell 2019, 176:535–548 e524.

15. Zeng T, Li YI: Predicting RNA splicing from DNA sequence using Pangolin. Genome Biol 2022, 23:103.

16. You N, Liu C, Gu Y, Wang R, Jia H, Zhang T, Jiang S, Shi J, Chen M, Guan MX, et al: SpliceTransformer predicts tissue-specific splicing linked to human diseases. Nat Commun 2024, 15:9129.

17. Chao KH, Mao A, Liu A, Salzberg SL, Pertea M: OpenSpliceAI provides an efficient modular implementation of SpliceAI enabling easy retraining across nonhuman species. Elife 2025, 14.

18. Radrizzani S, Kudla G, Izsvak Z, Hurst LD: Selection on synonymous sites: the unwanted transcript hypothesis. Nat Rev Genet 2024, 25:431–448.

19. Jang W, Park J, Chae H, Kim M: Comparison of In Silico Tools for Splice-Altering Variant Prediction Using Established Spliceogenic Variants: An End-User’s Point of View. Int J Genomics 2022, 2022:5265686.

20. Smith C, Kitzman JO: Benchmarking splice variant prediction algorithms using massively parallel splicing assays. Genome Biol 2023, 24:294.

21. Sorrentino BP, McDonagh KT, Woods D, Orlic D: Expression of retroviral vectors containing the human multidrug resistance 1 cDNA in hematopoietic cells of transplanted mice. Blood 1995, 86:491–501.

22. Tomberg K, Antunes L, Pan Y, Hepkema J, Garyfallos DA, Mahfouz A, Bradley A: Intronization enhances expression of S-protein and other transgenes challenged by cryptic splicing. bioRxiv 2021:2021.2009.2015.460454.

23. Ono Y, Doi N, Shindo M, Panico P, Salazar AM: Cryptic splicing events result in unexpected protein products from calpain-10 (CAPN10) cDNA. Biochim Biophys Acta Mol Cell Res 2022, 1869:119188.

24. Khan YA, Loughran G, Steckelberg AL, Brown K, Kiniry SJ, Stewart H, Baranov PV, Kieft JS, Firth AE, Atkins JF: Evaluating ribosomal frameshifting in CCR5 mRNA decoding. Nature 2022, 604:E16–E23.

25. Ansseau E, Domire JS, Wallace LM, Eidahl JO, Guckes SM, Giesige CR, Pyne NK, Belayew A, Harper SQ: Aberrant splicing in transgenes containing introns, exons, and V5 epitopes: lessons from developing an FSHD mouse model expressing a D4Z4 repeat with flanking genomic sequences. PLoS One 2015, 10:e0118813.

26. Baranick BT, Lemp NA, Nagashima J, Hiraoka K, Kasahara N, Logg CR: Splicing mediates the activity of four putative cellular internal ribosome entry sites. Proc Natl Acad Sci U S A 2008, 105:4733–4738.

27. Bosma PT, van Eert SJ, Jaspers NG, Stoter G, Nooter K: Functional cloning of drug resistance genes from retroviral cDNA libraries. Biochem Biophys Res Commun 2003, 309:605–611.

28. Cheng Y, Kang XZ, Chan P, Ye ZW, Chan CP, Jin DY: Aberrant splicing events caused by insertion of genes of interest into expression vectors. Int J Biol Sci 2022, 18:4914–4931.

29. Zeglinski K, Montellese C, Ritchie ME, Alhamdoosh M, Vonarburg C, Bowden R, Jordi M, Gouil Q, Aeschimann F, Hsu A: An optimized protocol for quality control of gene therapy vectors using nanopore direct RNA sequencing. Genome Res 2024, 34:1966–1975.

30. Lee JT, Yu SS, Kim VN, Kim S: Control of splicing efficiency by the mouse histone H2a element in a murine leukemia virus-based retroviral vector. Mol Ther 2007, 15:167–172.

31. Lemp NA, Hiraoka K, Kasahara N, Logg CR: Cryptic transcripts from a ubiquitous plasmid origin of replication confound tests for cis-regulatory function. Nucleic Acids Res 2012, 40:7280–7290.

32. Saffran HA, Smiley JR: The XIAP IRES activates 3’ cistron expression by inducing production of monocistronic mRNA in the betagal/CAT bicistronic reporter system. RNA 2009, 15:1980–1985.

33. Kowarz E, Krutzke L, Kulp M, Streb P, Larghero P, Reis J, Bracharz S, Engler T, Kochanek S, Marschalek R: Vaccine-induced COVID-19 mimicry syndrome. Elife 2022, 11.

34. Matsuoka K, Imahashi N, Ohno M, Ode H, Nakata Y, Kubota M, Sugimoto A, Imahashi M, Yokomaku Y, Iwatani Y: SARS-CoV-2 accessory protein ORF8 is secreted extracellularly as a glycoprotein homodimer. J Biol Chem 2022, 298:101724.

35. Ahmad M, Bellido Molias F, Cano Aroca L, Mordstein C, Watson S, Bhuiyan N, Clarke MJ, Kimchi-Sarfaty C, Katneni U, Gaunt E, et al: Splice-Aware Optimization Prevents Pervasive Missplicing of Natural and Synthetic cDNAs. bioRxiv 2026:2026.2005.2025.727620.

36. Burset M, Seledtsov IA, Solovyev VV: Analysis of canonical and non-canonical splice sites in mammalian genomes. Nucleic Acids Res 2000, 28:4364–4375.

37. Quarantani G, Clarke J, Thompson M, Sang F, Valcárcel J, Lehner B: OpenSplice: the impact of half a million mutations on the alternative splicing of 600 human exons. bioRxiv 2026:2026.2005.2022.727141.

38. Rowlands C, Thomas HB, Lord J, Wai HA, Arno G, Beaman G, Sergouniotis P, Gomes-Silva B, Campbell C, Gossan N, et al: Comparison of in silico strategies to prioritize rare genomic variants impacting RNA splicing for the diagnosis of genomic disorders. Sci Rep 2021, 11:20607.

39. You N, Liu C, Lin H, Wu S, Chen G, Shen N: Benchmarking pre-trained genomic language models for RNA sequence-related predictive applications. Nat Commun 2025, 17:223.

40. Xu C, Bao S, Wang Y, Li W, Chen H, Shen Y, Jiang T, Zhang C: Reference-informed prediction of alternative splicing and splicing-altering mutations from sequences. Genome Res 2024, 34:1052–1065.

41. Danis D, Jacobsen JOB, Carmody LC, Gargano MA, McMurry JA, Hegde A, Haendel MA, Valentini G, Smedley D, Robinson PN: Interpretable prioritization of splice variants in diagnostic next-generation sequencing. Am J Hum Genet 2021, 108:1564–1577.

42. Lord J, Oquendo CJ, Wai HA, Douglas AGL, Bunyan DJ, Wang Y, Hu Z, Zeng Z, Danis D, Katsonis P, et al: Predicting the impact of rare variants on RNA splicing in CAGI6. Hum Genet 2025, 144:243–251.

43. Rafi AM, Kiyota B, Yachie N, de Boer C: Characterizing homology-induced data leakage and memorization in genome-trained sequence models. bioRxiv 2026:2025.2001.2022.634321.

44. Anderson R, Ausler C, Jain A: When RNA goes off script: ensuring transcript fidelity in transgene expression. EMBO J 2026, 45:2420–2432.

45. Dao K, Jungers CF, Djuranovic S, Mustoe AM: U-rich elements drive pervasive cryptic splicing in 3’ UTR massively parallel reporter assays. Nat Commun 2025, 16:6844.

46. Chen XD, Jim M, Vallurupalli M, Cao K, Torres AN, Leong JW, Zhang Y, Wollensak D, Gong Q, Sun J, et al: Generative Design of Cell Type-Specific RNA Splicing Elements for Programmable Gene Regulation. bioRxiv 2025.

